# “Nanoscale biodegradable printing for designed tuneability of vaccine delivery kinetics”

**DOI:** 10.1101/2024.10.02.616252

**Authors:** David J Peeler, Rujie Sun, Ceren Kütahya, Patrick Peschke, Kun Zhou, Giulia Brachi, Jonathan Yeow, Omar Rifaie-Graham, Jonathan P Wojciechowski, Thomas Fernandez Debets, Vernon LaLone, Xin Song, Krunal Polra, John S Tregoning, Robin J Shattock, Molly M Stevens

## Abstract

Two photon polymerization (2PP) 3D printing enables top-down biomaterial synthesis with nanoscale spatial resolution for *de novo* design of monodisperse injectable drug delivery systems. To address the limitations of current 2PP resins, we developed Spatiotemporal Controlled Release Inks of Biocompatible polyEsters (SCRIBE), a novel poly(lactic-*co*-glycolic acid)-triacrylate resin family with sub-micron resolution and tuneable hydrolysis. SCRIBE enables direct printing of hollow microparticles with tuneable chemistry and complex geometries inaccessible to molding techniques, which we use to engineer controlled protein release *in vitro* and *in vivo*. We use SCRIBE microparticles to modulate antibody titers and class switching as a function of antigen release rate and extend these findings to enable a single-injection vaccine formulation with extended antibody induction kinetics. Demonstrating how the chemistry and CAD of 2PP-printed microparticles can be used to tune responses to biomacromolecule release *in vivo* opens significant opportunities for a new generation of drug delivery vehicles.

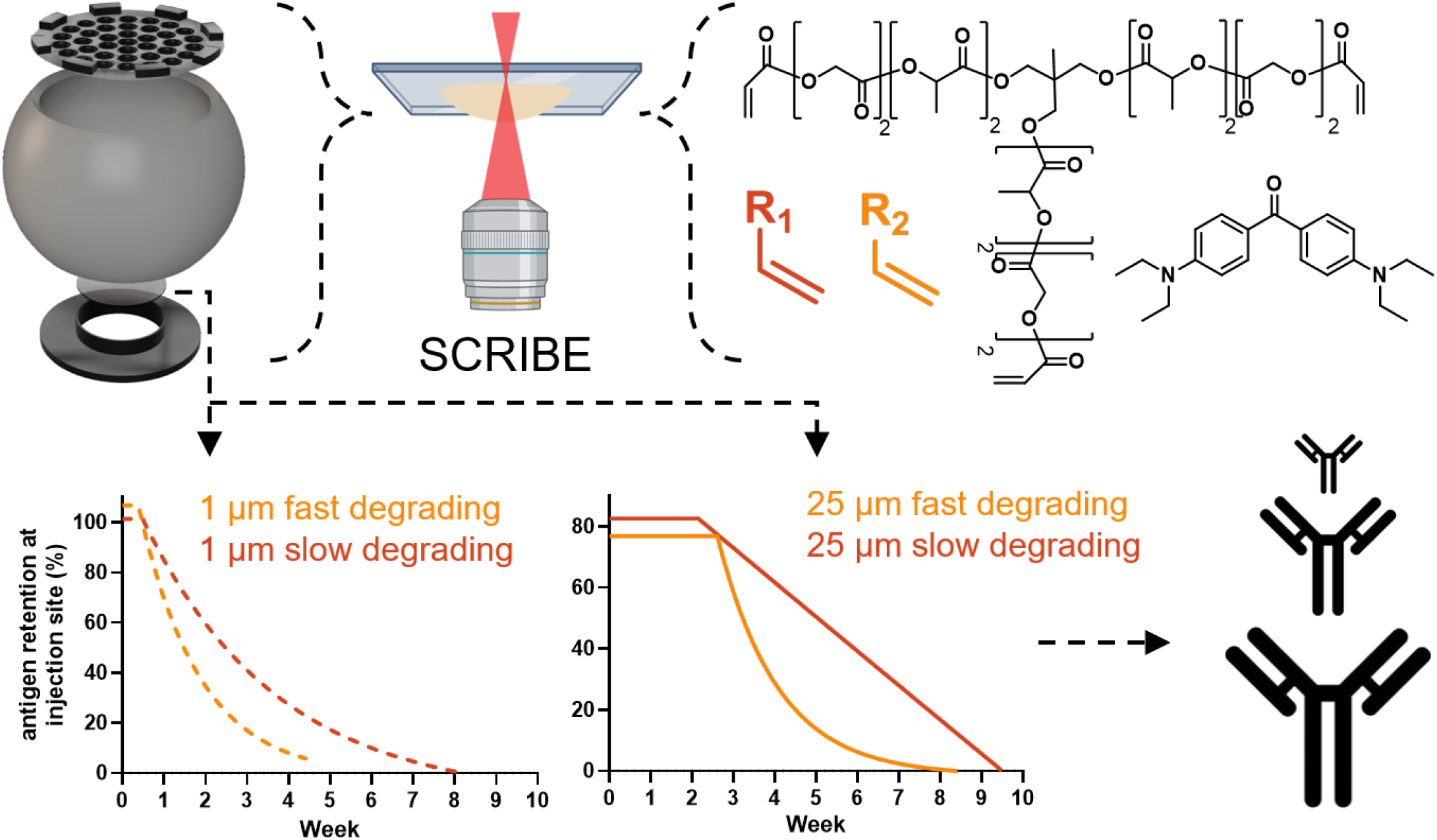

## Introduction

Microfabricated drug delivery systems have unique advantages over nanoscale formulations, including injection site retention, solvent-free loading, and integration into physically actuatable microrobotic systems. In recent years, hollow microparticles^1–4^ and microneedles^5^ that are sequentially loaded with cargo and sealed after fabrication have emerged as attractive delivery platforms based on their modularity and compatibility with various sensitive biomacromolecule cargos. Developed systems have primarily relied on moulding techniques that are restricted to simple geometries and lack broadly tuneable release profiles.^3–7^ Hollow delivery systems can achieve delayed, pulsatile release of vaccine booster doses without sacrificing antigen stability, enabling single-injection vaccination that could improve global health through increased compliance.^7–9^ Yet, platforms that allow for controlled, sustained vaccine delivery offer opportunities to further tune vaccine immunogenicity through persistent activation of lymph node germinal centers by sustained antigen release.^10–14^ Thus, a high-precision microfabrication method that can be easily chemically and spatially tuned could combine the advantages of injectable hollow particle platforms with the ability to engineer immunogenicity by tuning release rates.

We hypothesized that two photon polymerization (2PP) 3D-printing with biodegradable inks could enable top-down specification of delivery system geometry on length scales that are relevant for chemically-tuneable vaccine release. In recent years, 2PP has enabled sub-micron resolution printing with diverse materials including liquid crystal elastomers,^15^ silica,^16^ protein,^17^ and derivatives of classical biomaterials,^18–25^ yet none have been applied for drug delivery *in vivo*. We began by engineering hollow microparticles with commercially available, non-degradable 2PP inks to control drug release with stimuli-responsive sealants.^26–28^ However, for all examples of biomaterial 2PP, control of release has been limited to the intrinsic properties of the primary monomer used and printing throughput restricted by reliance on slow-scanning 63X objectives. Given recent demonstrations of high throughput micron-resolution printing with non-degradable resin,^2^ there remains a need for biocompatible, fast-polymerizing, high-resolution resins with tuneable degradation kinetics.

To address these shortcomings, we developed Spatiotemporal Controlled Release Inks of Biocompatible polyEsters (SCRIBE) and engineered a delivery system that demonstrates principles of both chemistry-and CAD-controlled release in mice. Optimization of poly(lactic acid-*co*-glycolic acid) (PLGA) macromonomer branching, molecular weight, and functionalization enabled copolymerization with small molecule comonomers at spatial resolutions on par with commercially available 2PP resins using an optical setup with 10-fold faster scan speed than previous 2PP biomaterial approaches. We further show that SCRIBE resin swelling and hydrolysis can be accelerated or decelerated through the selection of appropriate comonomers. Rapid prototyping on the Nanoscribe GT Photonic 2PP 3D-printer enabled the refinement of a hollow microparticle delivery system compatible with solvent-free protein loading and long-term encapsulation. Finally, we use SCRIBE to demonstrate that both CAD and comonomer chemistry can tune release of fluorescently labelled ovalbumin *in vitro* and *in vivo* and leverage this control to enable single-injection vaccination on par with prime-boost regimes. To our knowledge, this represents the first example of controlled release from biodegradable 2PP-printed systems *in vivo*. Building from a classic foundation in degradable PLGA, we illuminate a biocompatible strategy towards top-down engineering of chemically diverse biomaterials for drug delivery and beyond *via* two photon copolymerization.

## Main

### 1. PLGA-triacrylate enables sub-micron resolution printing with tuneable degradation

Resins for high resolution Dip-in-Laser-Lithography (DiLL) 2PP must be transparent for laser penetration, sufficiently viscous for substrate inversion/lens immersion, and rich in vinyl content for polymerization propagation. Building on previous work with branched poly(caprolactone)^19,21,22^ and poly(ethylene glycol)^23^ triacrylate 2PP resins, we designed a branched poly(lactic-*co*-glycolic acid)-triacrylate (PLGA-TA) macromonomer to serve as the biodegradable bulk of our 2PP resin, which was afforded at high yield *via* ring-opening polymerization followed by acrylation and thorough purification (**Fig. 1a-b; Supplemental Fig. 1-3**). Low molecular weight 1.2 kDa PLGA-TA had optimal solubility and viscosity compared to higher molecular weight macromonomers. Acrylating commercially available ε-caprolactone-trimethylolpropane (M_n_ = 830 Da; BOC Sciences, USA) yielded a poly(caprolactone)-triacrylate (PCL-TA) macromonomer for benchmarking (**Supplemental Fig. 4**). We formulated resin by solubilizing macromonomer and photoinitiator (EMK, 4,4’-bis(diethylamino)benzophenone) in the biocompatible precursor NVP (N-vinylpyrrolidone) to maximize resin vinyl content while simultaneously increasing print flexibility and hygroscopicity (**Fig. 1a**). In contrast to past work using the 63X Nanoscribe lens that maximizes light intensity and printing resolution,^19,21,22,25^ we engineered our resin to achieve the best resolution possible with the 25X lens at ten times faster printing speed (10^5^ vs 10^4^ mm/s). Using a resin formulation biased towards rapid polymerization (19% NVP, 1% EMK w/w), PCL-TA printing resolution was poor even at high laser power (90% lp) while PLGA-TA printing resolution was optimal at low laser power (30% lp), indicating a major difference in macromonomer reactivity (**Supplemental Fig. 5**). To maximize the advantage of PLGA biocompatibility, we reduced EMK and NVP content to 1% and 9% w/w, respectively, and re-optimized laser power. We observed high fidelity printing of resolution-benchmarking grid designs matching the commercially available, non-degradable IP-S resin (Nanoscribe GmbH, DE) at the maximum resolution achievable by the 25X lens used (XY = 500 nm, Z = 1 µm; **Fig. 1c-d; Supplemental Fig. 6**). We further hypothesized that varying comonomer content would impact PLGA degradation rate by modulating print hydration, pH, and flexibility. Acrylate monomers with a wide variety of side chain functionalities (carboxylic acids [CEA], tertiary amines [DMAEA], ethers [PEGA], alcohols [HEA], and crosslinkers [DEGDA]) were substituted for or blended with NVP to a final content of 9% w/w and used to print resolution benchmarking grids (**Supplemental Fig. 6**). All resins printed 1 µm-wide structures with < 150 nm error: equivalent or better than IP-S (**Fig. 1c**). To demonstrate how small differences in comonomer composition influence resin degradation rate, we chose to compare the 9% w/w NVP resin (“NVP”) to resins containing 4.5% CEA + 4.5% w/w NVP (“CEA”) and 4.5% DEGDA + 4.5% w/w NVP (“DEGDA”) throughout the rest of this work (**Fig. 1d; Supplemental Fig. 7**). Thus, formulation of a low molecular weight PLGA-triacrylate macromonomer with reactive comonomer(s) and a suitable photoinitiator enables high resolution 2PP with SCRIBE.

**Figure 1:**
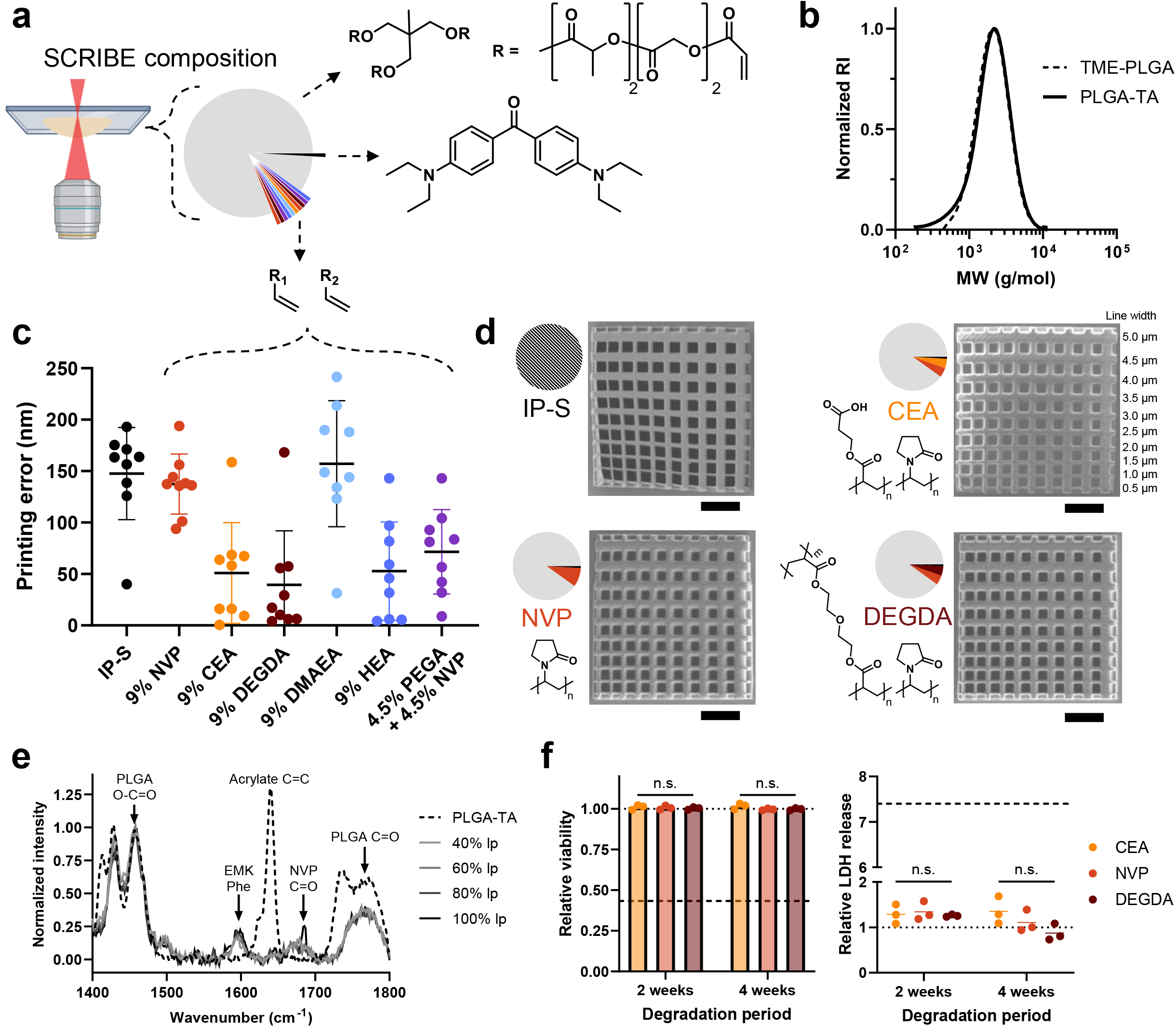
Spatiotemporal Controlled Release Inks of Biocompatible polyEsters (SCRIBE). (a) Illustration of SCRIBE resin raction (w/w): 90% PLGA-TA, 9% comonomer(s), 1% photoinitiator. (b) GPC characterization of TME-PLGA nd PLGA-TA macromonomer molecular weight (MW) distribution. (c) Quantification of printing error of 1 µm-es printed with commercial (IP-S) or SCRIBE resin formulations with the comonomers listed (data shown as [n = 9 measurements per grid]). (d) SEM micrographs of resolution grids printed with IP-S and SCRIBE resins VP, 4.5% CEA + 4.5% NVP, and 4.5% DEGDA + 4.5% NVP as co-monomers (w/w; scale bar = 20 µm). (e) ocal Raman spectra of pure PLGA-TA and SCRIBE NVP resin printed with varying laser power (lp; [n = 3 nts per sample]). (f) Human skeletal muscle cell viability (left, Alamar Blue assay) and membrane ation (right, LDH assay) following exposure to resin degradation products generated through particle incubation 0 °C for 2 or 4 weeks (data shown as mean ± SD [N = 3 biological replicates]; statistics from two-way ANOVA). corresponds to 500 µM Triton X-100 and dotted line corresponds to no treatment control.

We employed spectroscopic techniques and *in vitro* biocompatibility testing to characterize the composition of SCRIBE materials and the effects of their degradation products on human cells. In particular, we sought to confirm whether unpolymerized acrylate groups might be present after printing and/or pose biocompatibility concerns.^29^ Non-degradable commercially available resins like IP-DIP and IP-S (Nanoscribe GmbH) vary in vinyl bond polymerization during printing as a function of their monomer rigidity and photoinitiator content, with IP-DIP requiring post-print ultraviolet curing to achieve even 50% vinyl conversion.^30,31^ In contrast, our confocal Raman spectroscopy characterization revealed 96.8 ± 0.4% C=C bond conversion across all laser powers investigated without post-curing (**Fig. 1e**). Quantitative attenuated total reflectance Fourier transform infrared spectroscopy (ATR-FTIR) analysis of conversion was partially confounded by overlapping pyrrolidinone and ester absorption bands but qualitatively showed no difference between freshly printed and post-cured samples (**Supplemental Fig. 8**). Thus, blending a large PLGA-TA macromonomer with small, reactive comonomers enables efficient radical propagation and high acrylate conversion, maximizing PLGA content in the resin while retaining the ability to tune chemical properties through comonomer choice. We empirically validated the biocompatibility of our primary candidate resins (NVP, CEA, DEGDA) by applying their concentrated degradation products to human skeletal muscle cells and assessing cell viability (Alamar Blue™ assay) and membrane lysis (LDH assay). As demonstrated in **Fig. 1f**, SCRIBE degradation products are biocompatible, even when applied in extremely high concentrations.

### 2. Engineering in high resolution unlocks fabrication of protein-loaded hollow microparticles

Equipped with a biocompatible, tuneable 2PP resin, we engineered an innovative hollow microparticle design to evaluate macromolecule transport across SCRIBE materials. Rapid prototyping enabled by direct laser writing of SCRIBE particles enabled us to extensively optimize designs inaccessible to molding-based techniques (**Supplemental Fig. 9**). We aimed for injectability through 21-gauge needles or smaller and CAD-controlled cargo release through printed resin (as opposed to sealant-controlled release^26–28^). The evolved microparticle anatomy enables top-loading and bottom-release: protrusion, pores, chamber, porthole, and base (**Fig. 2a**). We have previously optimized lid pore size and protrusion height to balance chamber loading and dip-sealing.^26,27^ In this work, we optimized particle size to balance loading efficiency and injectability, settling on a 310 µm-wide sphere with sturdy 25 µm-thick walls and flattened top and bottom surfaces (volume = 8.5 nL). This geometry enables particle adhesion to the substrate during all fabrication steps and results in a nearly spherical particle that is readily injectable through a 21-gauge needle (**Fig. 2e-f**). Reducing particle diameter to 250 µm (volume = 4.3 nL; “Gen V”) enabled facile injection through 23-gauge needles but reduced cargo loading by ∼60%. Thus, we opted to use a smaller number of larger particles in our studies for simplicity. Implementation of automated loading procedures could unlock routine use of smaller particles as in other work.^4^ Inspired by osmotic pump delivery systems,^32^ we incorporated a “porthole” to serve as a release rate-controlling membrane and located this on the particle bottom to avoid mechanical stress and contact with the sealant. We hypothesized that porthole membrane thicknesses less than the wall thickness (25 µm) would enable us to evaluate whether SCRIBE could be used to control release at dimensions inaccessible to other fabrication approaches. Importantly, SCRIBE enables implementation of a round geometry and intricate lid and porthole features with sub-micron resolution impossible to replicate in a molding approach (**Fig. 2a**) and is also compatible with batch production (**Fig. 2b**) that may be adapted to scalable printing approaches.^2^

**Figure 2:**
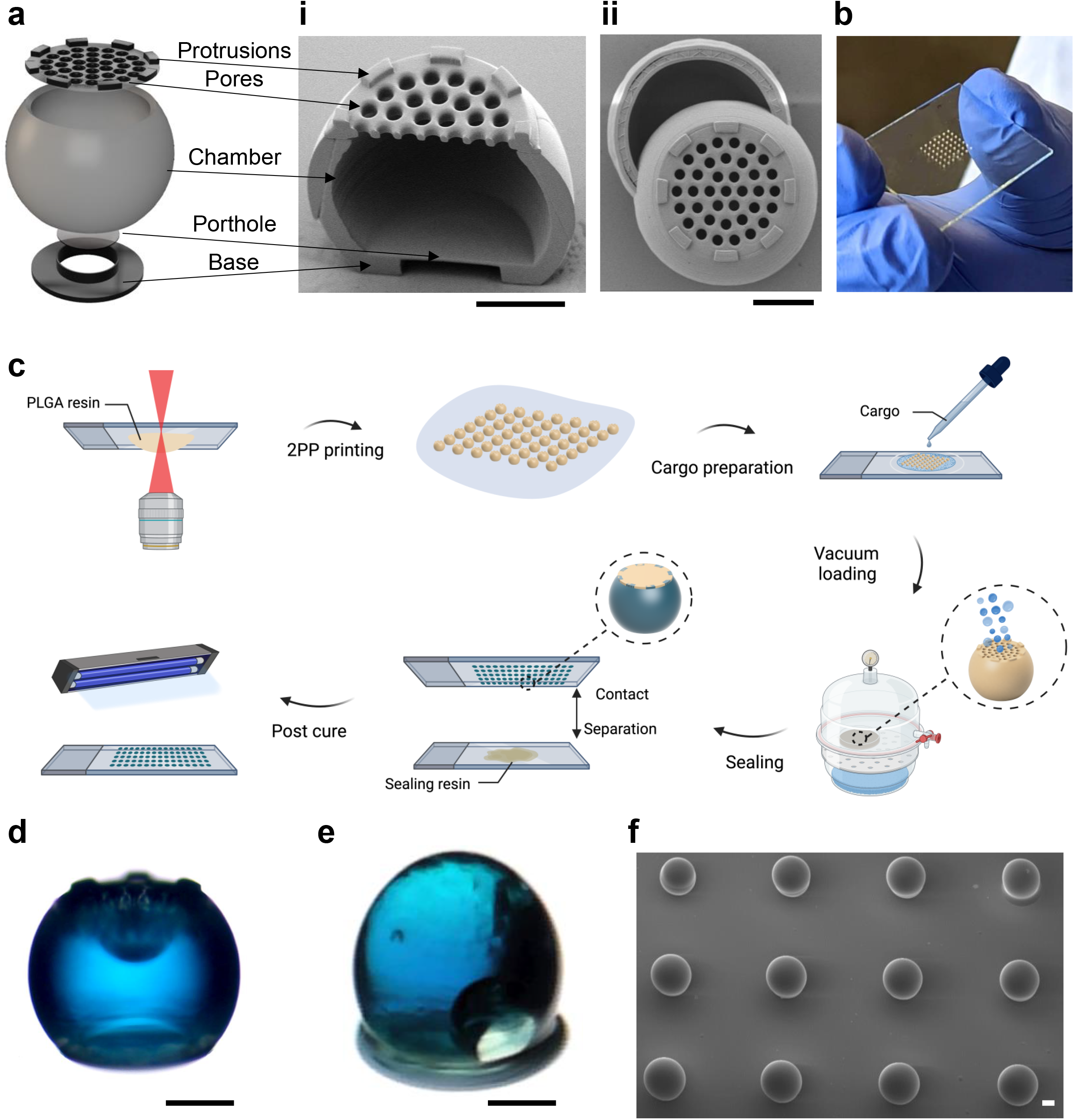
Hollow microparticle design and microfabrication. (a) Anatomy of an optimized hollow microparticle with porthole-elease with SEM micrographs of (i) XZ- and (ii) XY-plane cutaways. (b) A handheld 49 particle batch for . (c) Microfabrication process overview. (d) Brightfield image of a single particle after CF647-OVA loading and d (e) after sealing. (f) SEM of sealed particles ready for detachment and injection. All scale bars = 100 µm.

We adapted microfabrication techniques developed by our group ^26,27^ and others ^1–4^ to optimize the loading and sealing process shown in **Fig. 2c**. After printing and washing, a 20% w/w ovalbumin (OVA) solution in 1:1 v/v glycerol:water was loaded into particle batches under vacuum. Excess protein solution was recycled, and the particle exterior was cleaned with distilled water without disturbing the cargo chamber (**Fig. 2d**). To achieve long term encapsulation and evaluate printed resin-controlled release, we sealed the particles with a variant of the DEGDA SCRIBE resin with increased crosslinker content (25% DEGDA, 0% w/w NVP) and a UV-responsive photoinitiator (0.5% w/w Irgacure 369). The sealing resin was briefly brought into contact with the lid of the microparticles using an automated program on the rheometer and then cured with UV light. Lid pores prevent sealing resin from flowing into the chamber due to surface tension and the protrusions control the sealing layer thickness. Particles loaded with fluorescent OVA (CF647-OVA) were visibly blue when imaged by light microscopy (**Fig. 2e**) and displayed a smooth, spherical shape as shown by SEM (**Fig. 2f**). Differences in protein loading between particles of different resin chemistries were statistically insignificant, although batch-to-batch loading was less variable for DEGDA particles (CEA = 516 ± 150; NVP = 596 ± 135; DEGDA = 435 ± 52 ng OVA/particle determined by microBCA; **Supplemental Fig. 10**). This loading is comparable to that achieved by others with similar sized particles, but with 50-fold thinner particle wall thickness.^1–4^ Thus, we applied SCRIBE to establish a purely monodisperse injectable delivery platform enabling the first investigation of how single micron-thick biodegradable materials control macromolecule release.

### 3. SCRIBE comonomer choice controls particle swelling and release *in vitro*

In comparison to particle molding approaches that rely on solvent-casted blends of miscible polymers^3,4^ or 2PP approaches using protein^17^ or GelMA^25^ hydrogels with large polymer network voids, the chemical diversity of small monomers compatible with SCRIBE offers vast opportunity for tuneable release. We observed that employing ionizable comonomers (CEA, DMAEA) at 10 wt% dramatically accelerated SCRIBE print swelling and degradation at 50 °C in PBS compared to neutral monomers (NVP, HEA, PEGA), leading to total degradation in less than two weeks (**Supplemental Fig. 11**). In contrast, substituting the non-degradable crosslinker DEGDA decreased, but did not eliminate print swelling and IP-S particles exhibited no swelling as expected. Scanning electron microscopy of 1 µm-porthole CEA particles (4.5% CEA, 4.5% w/w NVP) incubated at 37 °C in PBS revealed that portholes were deformed as early as day 5, ruptured by day 25, and fully eroded or detached by day 30 (**Fig. 3a**). Degradation of CEA resin revealed the interior scaffold pattern of the particle and proceeded to complete disintegration by week 6, consistent with bulk erosion of PLGA. By comparison, NVP particles did not visibly deform until 3 weeks of incubation and DEGDA (4.5% DEGDA, 4.5% w/w NVP) particles exhibited minimal deformation over the course of 10 weeks of incubation. Thus, comonomer choice dictates SCRIBE resin hydration and degradation rate, even when blended 1:1 w/w with NVP and printed at equivalent laser power.

**Figure 3:**
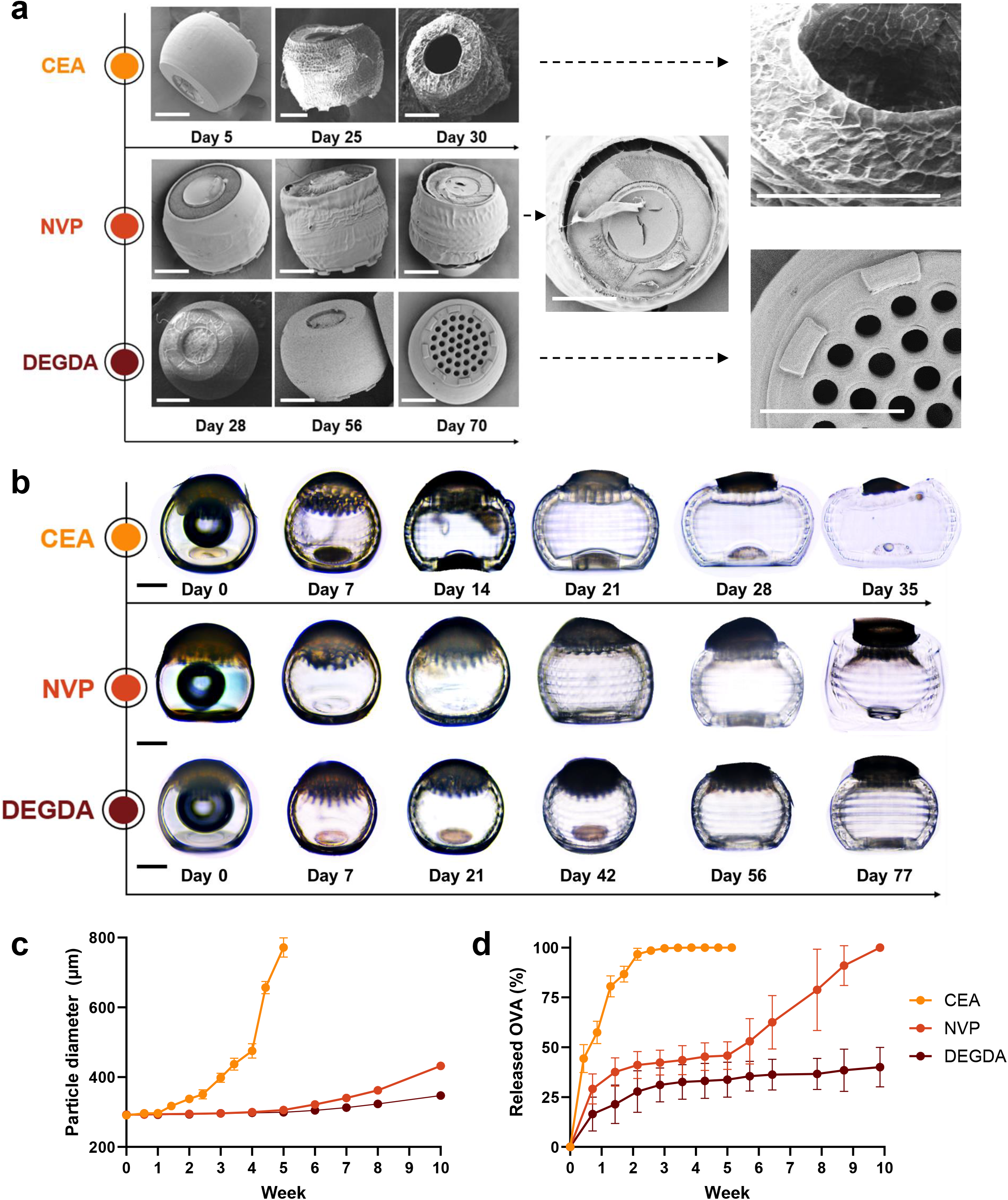
Comonomer choice controls particle swelling and release. (a) SEM and (b) light microscopy of SCRIBE particles comonomers (2-carboxyethyl-acrylate [CEA], N-vinylpyrrolidone [NVP], diethyleneglycol diacrylate [DEGDA]) PBS at 37 °C. (c) Swelling of microparticle diameter as quantified from light microscopy images (data shown SD [n = 10 particles]). (d) Ovalbumin release from 1 µm-porthole particles incubated in PBS at 37 °C as y microBCA (N = 4 batches). Because full degradation did not occur in the timeline studied, DEGDA release sed to the average loading of separately measured batches. All scale bars = 100 µm.

We next fabricated OVA-loaded, 1 µm-thick porthole particles with various blended resin chemistries and monitored protein release and particle swelling during incubation at 37 °C in PBS over 10 weeks. Sealed particle swelling followed similar trends to the unsealed particles evaluated by SEM, with CEA, NVP, and DEGDA particles reaching 1.25X their initial diameter at days 10, 39, and 50, respectively (**Fig. 3b-c**). All 1 µm particles released OVA with first order kinetics over the first two weeks of incubation, corresponding to 100%, 33%, and 25% of the total encapsulated OVA for CEA, NVP, and DEGDA particles, respectively (**Fig. 3d**). NVP particles exhibited further zero order release kinetics from week 5-8 while DEGDA particles did not release any further cargo. The biphasic release observed for the NVP-only resin may imply that protein-particle interactions may restrict exit until sufficient PLGA degradation has occurred. By contrast, 25 µm porthole particles did not exhibit release in the time frames studied, which may be attributed to the limit of detection of the assay used as we have confirmed that long term (1 month) OVA encapsulation in slow releasing particles (25 µm-thick NVP) does not result in degradation or aggregation (**Supplemental Fig. 12**). These results demonstrate that SCRIBE polymer network density allows for immediate transport across 1 µm porthole thicknesses according to Fick’s laws of diffusion. Importantly, SCRIBE release rate can be modulated by weeks simply by selecting comonomers to alter resin swelling and degradation.

### 4. Design- and chemistry-controlled release shapes immunogenicity *in vivo*

Having observed weeks-long sustained OVA release *in vitro*, we hypothesized that SCRIBE could offer the opportunity to tune vaccine immunogenicity through sustained vaccine delivery. We monitored the subcutaneous release of 20 µg CF647-OVA from 1 and 25 µm porthole particles printed with CEA and NVP using *in vivo* fluorescent imaging and simultaneously monitored serum antibody concentrations. Soluble CF647-OVA (± *ad hoc* mixed empty CEA particles) was >95% cleared within 24 hours, implying that fluorescent signal at the injection site at later timepoints in other groups could be attributed to particle-encapsulated CF647-OVA (**Fig. 4a**). The fluorescence intensity of encapsulated CF647-OVA reached a maximum at day 1-3 for 1 µm-thick particles and day 1 (CEA) or day 7 (NVP) for 25 µm-thick particles, reflecting *in vitro* data implying that resin hydrophilicity and thickness regulate diffusion of water across the particle wall (**Fig. 4b-c**). We observed agreement between OVA release kinetics *in vivo* and *in vitro* for 1 µm-thick CEA particles, but *in vivo* release from 1 µm-thick NVP particles was more rapid than observed *in vitro* and exhibited first order kinetics (**Fig. 4b**). In contrast to near-immediate release observed for 1 µm-thick particles, 25 µm-thick particles exhibited a delayed onset of release beginning at week 3 (**Fig. 4c**). Resin chemistry further dictated release rate from 3 weeks onward, with CEA particles exhibiting 3 weeks of first order release and NVP particles exhibiting 6 weeks of zero order release. Therefore, SCRIBE particle thickness and chemistry influenced the kinetics of vaccine release *in vivo*, with sustained delivery profiles ranging from weeks to months. In contrast to the delayed burst release of other hollow microparticle approaches,^3,4^ SCRIBE may offer the opportunity to tailor both the timing and kinetics of “extended priming” to synchronize with innate cell recruitment and germinal center initiation.^14^

**Figure 4:**
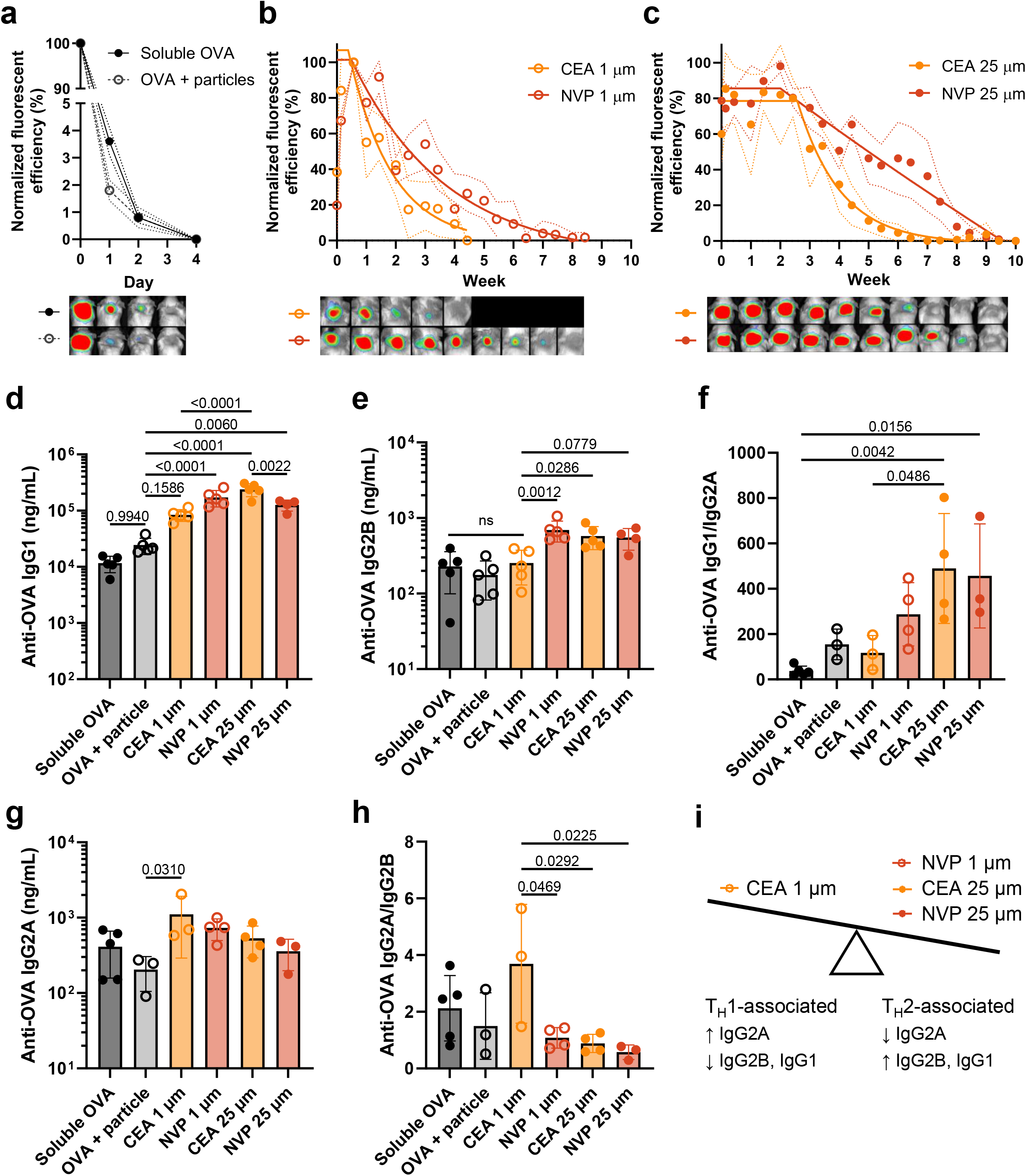
Design- and chemistry-controlled release shapes immunogenicity *in vivo*. Longitudinal *in vivo* fluorescent imaging mice injected subcutaneously with 20 µg CF647-OVA delivered in (a) soluble form ± empty CEA particles or ed in CEA or NVP particles with either (b) 1 µm or (c) 25 µm porthole thickness. Background-subtracted e ROI fluorescent efficiency (λ_ex_/λ_em_ = 640/690 nm) was normalized to the maximum observed per mouse (Data mean [circles] ± SD [dashed lines] with the plateau followed by one phase decay fit [solid line; N=3-4]; efficiency scale for raw images is 0.4-1*10^-4^ units/cm^2^). Anti-OVA (d) IgG1 titers, (e) IgG2B titers, (f) titer ratios, (g) IgG2A titers and (h) IgG2A/IgG2B titer ratios in week 8 serum plotted as mean ± SD (N=3-5; p-ne-way ANOVA with Tukey’s multiple comparisons test). (i) Illustration formulation-induced antibody bias.

We next sought to correlate differences in antigen release rate with humoral responses. As in other work,^13,33,34^ anti-OVA IgG responses were predominantly of the IgG1 subclass. All SCRIBE formulations achieved significantly greater anti-OVA IgG1 titers at week 8 compared to soluble OVA, but only the slowest-releasing formulations (1 µm NVP, 25 µm NVP, 25 µm CEA) achieved significance relative to OVA mixed with empty CEA particles (**Fig. 4d**). As in release assays, differences in IgG1 titers could be explained by both porthole thickness (1 vs 25 µm CEA, p < 0.0001) and chemistry (25 µm CEA vs NVP, p = 0.0022). Differences between 1 and 25 µm CEA particles are particularly interesting as both release antigen at similar rates (first order release lasting ∼3 weeks) but begin releasing at different times (3- or 21-days post-injection, respectively). Analysis of anti-OVA IgG2 subtypes revealed that slow-releasing formulations skewed towards T_H_2-associated humoral responses with higher IgG2B titers (**Fig. 4e**) and IgG1/IgG2A ratios (**Fig. 4f**) compared to soluble controls and fast releasing 1 µm CEA particles. Conversely, fast-releasing 1 µm CEA particles skewed towards T_H_1-associated humoral responses, achieving the highest IgG2A titers (**Fig. 4g**) and significantly higher IgG2A/IgG2B ratios compared to other particle formulations (**Fig. 4h**). Our results agree with others who have shown that sustained OVA release correlates with increased IgG1 titers compared to rapidly cleared antigen.^13,33,34^ Here, however, we leverage SCRIBE to identify a subtle effect threshold between immediate zero order release lasting three weeks (T_H_1-biased) and longer-term release profiles (T_H_2-biased; **Fig. 4i**).

### 5. SCRIBE enables single injection vaccination on par with prime-boost regimes

We then benchmarked single-injection SCRIBE vaccine formulations against a prime-boost regime to evaluate their potential for increasing compliance and protection in resource-constrained settings. We specifically aimed to understand whether the release rate efficacy threshold identified in the prime-only study remained relevant when particles were co-injected with soluble antigen. Thus, mice received a single injection containing a prime of 5 µg soluble OVA + 20 µg CF647-OVA encapsulated in either 1 µm CEA or 1 µm NVP particles to serve as boost. Control mice received a 5-µg soluble CF647-OVA prime and a boost injection of 20 µg soluble OVA on day 14; antigen release and anti-OVA IgG were monitored as before (**Fig. 5a**). Particle-mediated antigen release was immediate but sustained as before (NVP slower than CEA; **Supplemental Fig. 13**), resulting in different kinetics of anti-OVA IgG production compared to control mice receiving separate injections of fast-draining soluble antigen. Mean anti-OVA IgG increased by 49.1-fold from weeks 2-4 and 5.7-fold from weeks 4-8 in response to two soluble doses, whereas NVP-boosted titers rose more gradually (6.7- and 3.8-fold, respectively; **Fig. 5b**). Both particle-boosted groups exhibited higher mean anti-OVA IgG titers at week 2 compared to the control group before boosting. (**Fig. 5c**) Rapid boosting from 1 µm CEA particles resulted in mean anti-OVA IgG titers that were significantly lower than control at week 4 (p = 0.0388) and contracted from 46% from week 6-8, echoing results from the prime-only study. In contrast, sustained boosting from 1 µm NVP particles resulted in titers that were not statistically different from the double-injection control group at weeks 4-8. (**Fig. 5d**) Intriguingly, anti-OVA IgG1/IgG2A and IgG2A/IgG2B ratios were not significantly different between groups, indicating that particle-mediated subtype biasing may only be possible in the absence of a soluble priming dose (**Fig. 5e-f**; **Supplemental Fig. 14**). Thus, a single injection OVA vaccine formulated in SCRIBE particles with optimized thickness and degradation rate can achieve similar antibody titers compared to a two-injection soluble formulation, and with a more rapid onset of immunity. Finally, we observed that SCRIBE particles appear to be either fully degraded or highly biocompatible *in vivo*, as no particles or indicative signs of inflammation (e.g. granuloma, vascularization, swelling, edema, etc.) were observed in the subcutaneous neck scruff following sacrifice at day 61 (**Fig. 5g**; **Supplemental Fig. 15**) or in the subcutaneous flank following sacrifice at day 49 (**Supplemental Fig. 16**), highlighting the translational potential of SCRIBE materials.

**Figure 5:**
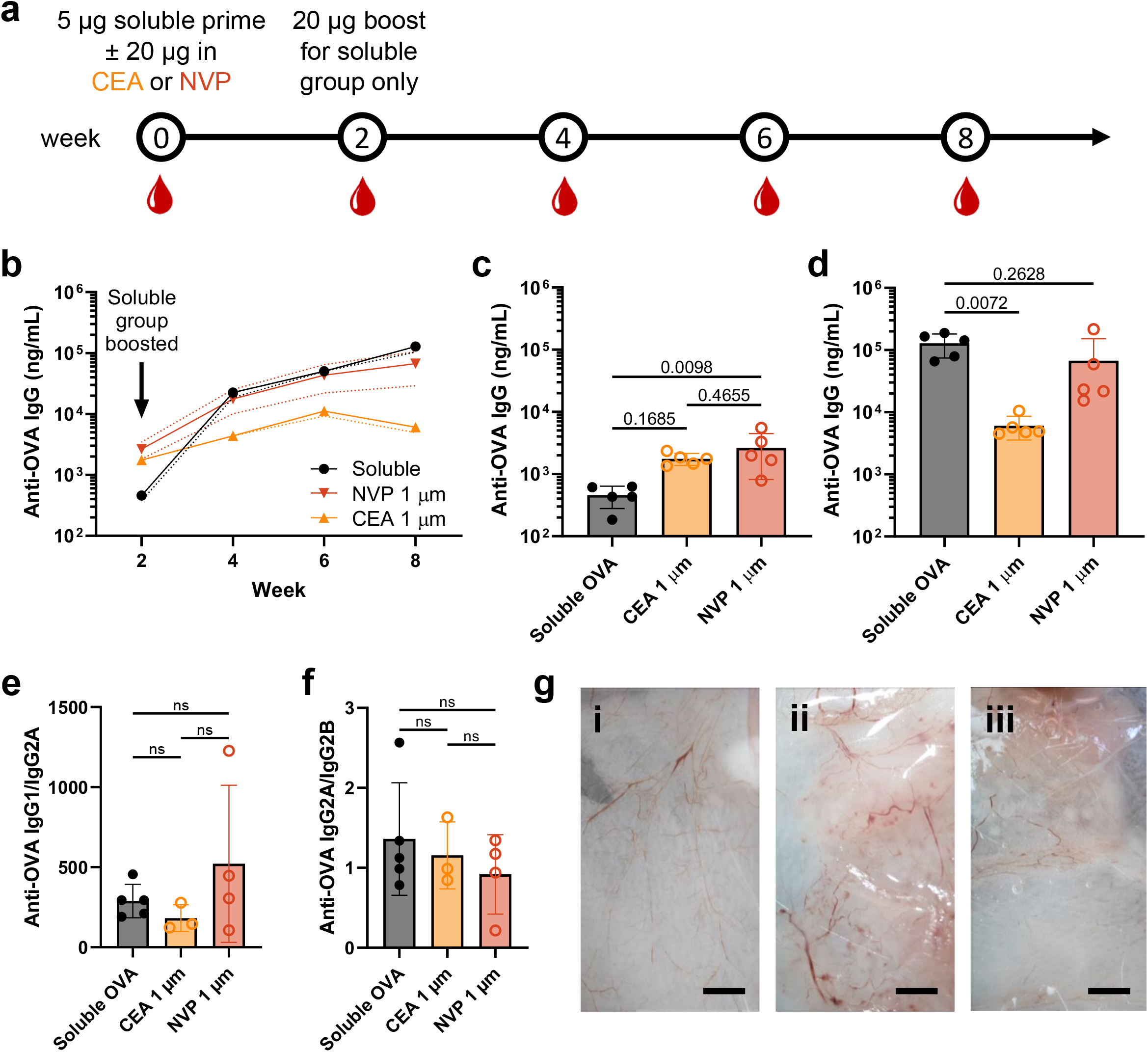
SCRIBE enables single injection vaccination on par with prime-boost regimes. (a) Prime-boost tion and serum bleeding schedule. (b) Time course, (c) week 2, and (d) week 8 of anti-OVA IgG serum ek 8 anti-OVA (e) IgG1/IgG2A and (f) IgG2A/IgG2B titer ratios. (Data shown as mean ± SD [N = 3-5], p-ia one-way ANOVA with Tukey’s multiple comparisons test.) (g) Photographs of the neck scruff eous layer dissected at day 61 from mice receiving antigen in (i) soluble form or encapsulated in (ii) CEA or particles. Scale bar = 4 mm.

## Conclusions and Outlook

We introduce SCRIBE through direct laser writing of PLGA microparticles for controlled protein release. Optimized SCRIBE resins allow for sub-micron resolution printing at significantly faster speeds than previous 2PP approaches while leveraging the inclusion of diverse comonomers to fine-tune print swelling and degradation independent of resolution. By engineering a sealable hollow microparticle, we were able to evaluate macromolecule transport across uniform micron-thick degradable biomaterials. We applied SCRIBE particles to deliver a model vaccine and found specific requirements for sustained release profiles that correlate with maximal antibody titers and subtype class switching. Ultimately, particles with 1 µm thickness printed with a relatively slow degrading SCRIBE resin could deliver a model vaccine booster dose with the right kinetics to induce equivalent titers to a double injection formulation. These results open the possibility to translate recent demonstrations of neutralizing antibody maturation via “extended priming” into single injection vaccine formulations.^14^

We did not scrutinize innate immune responses in this work as we did not observe any statistically significant differences in antibody titers or subtype ratios when comparing soluble OVA co-injected with empty CEA particles to soluble OVA injected alone. Furthermore, comparing results from 1 µm and 25 µm CEA particles (which differ only in porthole thickness) enables us to conclude that antigen release rate and not direct effects of material degradation result in different antibody titers and classes. Taken together, these results imply that SCRIBE particles do not provide the innate immune stimulus ascribed to other PLGA microparticles,^3^ opening the opportunity to tailor immune responses via adjuvants. Understanding the mechanisms of material- or adjuvant-driven innate stimulation and their intersection with optimal antigen delivery kinetics is a promising area of future research that SCRIBE offers great flexibility to investigate.^14^ We envision that SCRIBE will be the foundation of advanced microfabrication processes that implement emerging high throughput printing modalities and automated loading robots. With SCRIBE, we offer a future where top-down drug carrier designs are drawn in high resolution with biocompatible ink.

## Methods

### Materials

All chemicals were purchased from Sigma Aldrich and used without further purification unless otherwise specified. Other materials were purchased from the suppliers indicated in the text.

### Conjugation of CF647-OVA

Albumin from chicken egg white (OVA; Sigma Aldrich) was dissolved at 40 mg/mL in 0.1 M sodium bicarbonate (213.5 mg; 5 µmol) and placed under magnetic stirring at room temperature. An equivalent molar quantity of CF™ 647 succinimidyl ester (CF647; Sigma Aldrich) was dissolved at 10 mM in anhydrous DMSO, added dropwise into the stirred OVA solution, and allowed to react for 2 hours. The CF647-OVA product was purified by dialysis (Snakeskin 10 kDa cutoff; Sigma Aldrich) against water and lyophilized for future use.

### PLGA-TA macromonomer synthesis, purification, characterization

#### a. Synthesis and purification

##### TME-PLGA precursor synthesis

D,L-lactide and glycolide were recrystallized in toluene prior to use in the synthesis of precursor (TME-PLGA). Monomer ratios of 1:1 (D,L-lactide/glycolide) were used in the synthesis, targeting a molar mass of 1 kDa. For the polymerization, lactide (5.99 g, 41.6 mmol), glycolide (4.83 g, 41.6 mmol), and 1,1,1-tris(hydroxymethyl)ethane (TME; 1.50 g, 12.5 mmol) were brought to room temperature, weighed, and transferred to a flame-dried flask equipped with a reflux condenser under a gentle nitrogen flow. The flask was sealed, and the chemicals were dried under vacuum for at least 3 hours. The reaction was then conducted in bulk under a nitrogen atmosphere, with the monomers and initiator heated to 130 °C. Once all the reactants had melted, tin(II) 2-ethylhexanoate (26.9 µL, 0.08 mmol) was added as catalyst. The polymerization was terminated after 2 hours by cooling after which the reaction was diluted in DCM, and the resulting polymer was precipitated in a cold 1:1 v/v mixture of hexane and diethyl ether and subsequently dried under air flow.

##### PLGA-TA macromonomer synthesis

Approximately 10 g of TME-PLGA was dissolved in 50 mL of anhydrous DCM. An equal mass of potassium carbonate was then added relative to the mass of the polymer, resulting in the formation of a slurry. The reaction flask was then cooled using an ice/salt bath before distilled acryloyl chloride was added at a 1.5 molar ratio relative to the PLGA hydroxyl groups dropwise under continuous stirring. The reaction was allowed to stir at room temperature for two days. Upon completion of the reaction, the mixture was filtered under vacuum to remove the solids and subsequently washed three times with DCM. The resulting solution was concentrated, yielding functionalized PLGA as a viscous liquid. The resultant polymer was precipitated in a cold 1:1 v/v mixture of hexane and diethyl ether and dried under air flow. Subsequently, 5 g of functionalized polymer was dissolved in DCM and dialyzed in 1:1 v/v mixture of DCM and THF using a Pur-A-Lyzer™ Mega Dialysis Kit (MWCO 1 kDa) for 3 days. The polymer was then precipitated in cold 1:1 v/v mixture of hexane and diethyl ether and dried under air flow.

##### PCL-TA macromonomer synthesis

Commercially available ε-Caprolactone-trimethylolpropane (M_n_ = 830 Da; BOC Sciences, USA) was acrylated using the same procedure described for PLGA-TME above. Due to incompatibility with ether precipitation, purification was instead performed by silica column chromatography as previously described for PCL-trimethacrylate.^35^

#### b. ^1^H and ^13^C NMR Characterization

^1^H and ^13^C NMR spectra were acquired on a Bruker Avance 500 MHz NMR spectrometer operating at 293 K. Deuterated chloroform (CDCl_3_) was acquired from Sigma Aldrich and used as received. Chemical shifts (δ) were referenced to the residual solvent peak (δ = 7.26 ppm). Proton (^1^H) NMR data are reported as chemical shifts with the following multiplicity notation: s = singlet; d = doublet; t = triplet; q = quartet; m = multiplet; br = broad; td = triplet of doublets. This is followed by the proton position and then the coupling constants (*J*) in Hertz if applicable.

#### c. Gel Permeation Chromatography

Macromonomer and precursor molecular weight distributions were characterized using an Agilent PL GPC-50 instrument equipped with a refractive index (RI) detector running in HPLC grade DMF (containing 0.075 wt % LiBr) at a flow rate of 1.0 mL min^−1^ at 40 °C through two GRAM Linear columns (Polymer Standards Service) in series. Near monodisperse poly(methyl methacrylate) standards dissolved in eluent were used to calibrate the instrument. Polymer was dissolved in eluent at 2 mg/mL and filtered through a 0.2 μm syringe filters prior to analysis.

#### d. LC-MS

LC-MS spectra was acquired on a Shimadzu LC-2060C UHPLC coupled to a Shimadzu LCMS-2050 single quadrupole, dual source (ESI/APCI) mass spectrometer. The mass spectrometer used a nebulizing gas flow = 2.0 L/min, drying gas flow = 5 L/min, heating gas flow = 7 L/min, desolvation temperature = 450 °C and desolvation line temperature = 200 °C. The column was a 50 × 2.1 mm Phenomenex Kinetex EVO C18, particle size = 2.6 Cm, pore size = 100 Å run at a flow rate of 0.5 mL/min with an injection volume of 5 µL. Mobile phases were: A = HPLC grade water (VWR International) with 0.1% formic acid and B = HPLC grade acetonitrile (VWR International) with 0.1% formic acid. A gradient elution was performed using the method as follows: 0 – 1.5 min 5% B, 1.5 - 5.5 min 95% B, 5.5 - 7 min 95% B, 7 – 7.1 min 5% B, 7.1 - 12 min 5% B. In a typical experiment, PLGA-TA was dissolved at 10 µM in HPLC grade methanol and filtered through a 0.45 µm PTFE filter before injection. Positive mode ESI/APCI mass spectra were extracted from the centre of the polymer envelope (4.502 - 5.001 min).

### Resin formulation

Comonomers were either used as supplied (NVP, N-vinyl pyrrolidone) or first passed through a small column of inhibitor removing resin (CEA, 2-carboxy ethyl acrylate; DMAEA, 2-dimethyl amino ethyl acrylate; HEA, 2-hydroxy ethyl acrylate; PEGA, poly(ethylene glycol) methyl ether acrylate [M_n_ = 470]; DEGDA, diethylene glycol diacrylate) and stored at -20 °C for further use. The photoinitiator EMK (4,4’-bis(diethylamino)benzophenone) was dissolved at 10% w/w in each comonomer, which were then blended to achieve the desired comonomer composition. Printing resin was prepared by adding comonomer-photoinitiator to PLGA-TA for a final photoinitiator : comonomer : macromonomer weight ratio of 1 : 9 : 90, dissolved using approximately 0.5 vol eq dichloromethane, and stored at 4 °C in the dark for further use. Sealing resin was prepared at a final photoinitiator : comonomer : macromonomer weight ratio of 0.5 : 24.5 : 75 using Irgacure 369, DEGDA, and PLGA-TA, respectively. Dichloromethane was removed before printing or sealing by placing resin under vacuum after deposition on substrate.

### Two photon polymerization printing

A commercial 2PP printer (Photonic Professional GT2, Nanoscribe) was utilized to print hollow microparticles with intricate micro features using custom PLGA resins. The microparticles were designed using commercial computer-aided design software (Autodesk Inventor 2023), and the model files (in STL format) were exported. The model files were then processed in a job development software (Describe 2.7, Nanoscribe), where a shell printing strategy was adopted to balance reasonable printing times and maintain structural features with high mechanical strength. The shell printing technique uses outer shells to ensure structural features, while interior scaffolds support the whole structure, significantly reducing printing time. The optimized printing parameters were: fixed mode with Z slicing distance 1 µm; shell parameters with XY hatching distance 0.5 µm, contour count 12, and base slice count 2; triangle scaffold with XY hatching distance 0.5 µm; each particle printed within a single block. The laser power settings were optimized for each resin as described in the text.

ITO-coated glass was used as the substrate for printing the microparticle array. The ITO coating increases the refractive index contrast, enabling identification of the printing interface between resin and glass substrate. To further enhance adhesion, the substrate was activated by O2 plasma (GaLa Instrumente Prep 5, 0.3 bar, 150 W) for 5 minutes to make it hydrophilic. For printing, the treated ITO-coated glass was placed into its dedicated holder, and the resin was applied to cover a sufficient printing area. To prevent interaction between the laser and any tiny bubbles inside the PLGA, the resin on the substrate was left under vacuum for 1 minute to eliminate the bubbles. Using a 25× NA0.8 objective, the printing process typically lasted 2 hours for a batch array of 7×7 particles. After printing, uncured PLGA resin was removed by immersing the entire substrate in a beaker with PGMEA for one hour, followed by submersion in IPA for 10 minutes to remove excess PGMEA.

### Characterization of printed resin

#### a. Scanning electron microscopy and quantification of printing error

Samples were transferred to conductive tape on the top surface of a metal stub, either attached to glass substrates for freshly printed samples or directly on the tape for degradation study samples. Degradation study samples were washed three times with water to remove buffer salts, rapidly frozen in water at -80 °C, and lyophilized. Samples were either coated with 15 nm chromium in a pumped coater (Quorum, Q150T Plus) before scanning electron imaging was conducted on a Zeiss Auriga microscope with 5 kV voltage, or they were coated with gold (30 s deposition time, 20 mA current) via sputtering deposition (Emitech K575X peltier cooled) before scanning electron imaging was conducted on a Zeiss Leo Gemini with 3 kV accelerating voltage.

Printing error was quantified using ImageJ and Python (see Supplementary Information and Source Data for code used). SEM images (5000X magnification) of resolution grids were first uniformly contrast enhanced (1% saturated pixels; histogram normalized) and sharpened in ImageJ to normalize variation inherent in imaging. Pixel intensity plots traversing the smallest five lines were exported from ImageJ for each image. Jupyter notebook was used to run a custom Python script (“FWHM.ipynb”) to smooth line profiles with a rolling average and then measure the full width half maximum (FWHM) of each line. Printing error was determined as the absolute difference between specified and observed line width. Portions of the Python code used were developed with assistance from Claude 3.5 Sonnet, an AI assistant created by Anthropic, PBC.

#### b. Confocal Raman spectroscopy

Cubes (250 µm^3^) were printed with a solid fill program at varying laser power (40, 60, 80, 100%), washed with PGMEA and IPA, and allowed to dry. Prints were then submerged with distilled water and imaged with a WITec alpha 300R + Raman confocal microspectrometer equipped with a piezoelectric stage (UHTS 300, WITec GmbH), 63X water immersive objective lens (Zeiss W Plan Apochromat 63X, N.A = 1), red solid-state excitation laser (λ = 785 nm, 85 mW, WITec GmbH) and an imaging spectrograph (Newton, Andor Technology Ltd). This setup enabled acquisition of spectral data across a wavenumber range from 0-2600 cm^−1^. Five representative spectra were gathered in different locations of each sample, which were averaged, processed to remove cosmic rays, background subtracted, and normalized to signal at 1453 cm^-1^ (corresponding to PLGA ester signal; 1405.0 to 1485.0 cm^-1^) using ProjectFIVE software (WiTec). Spectra from pure PLGA-TA were collected and processed identically for comparison. Percent residual/unconverted vinyl was calculated by dividing the area under the curve for the peak at 1640 cm^-1^ (1627.5 to 1652.5 cm^-1^) for printed structures by that of pure PLGA-TA, with % conversion given as the complement (100% - % unconverted).

#### c. Fourier Transform Infrared Spectroscopy (FTIR)

Three 1 mm^3^ cubes were printed with a solid fill program on separate substrates at 80% laser power, washed with PGMEA and IPA, allowed to dry, and then subjected to either 0, 30, or 60 minutes of post-curing UV exposure under a 5 mW/cm^2^ 365 nm lamp. Prints were then subjected to AT-FTIR analysis on a Perkin Elmer Spectrum 100 spectrometer at a resolution of 1 cm^-1^ with 52 scans. Spectra were trimmed to 1550-1850 cm^-1^ to center on the fingerprint region of interest surrounding the acrylate C=C stretch (1635 cm^-1^), NVP C=O stretch (1675 cm^-1^), and ester C=O stretch (1750 cm^-1^) and normalized to the transmittance at 1750 cm^-1^.

### Degradation product biocompatibility

Three batches (49 particles/batch) printed with NVP, CEA, or DEGDA resin were separately incubated in 150 µL PBS at 50 °C for 2 weeks to promote accelerated degradation. Buffer was replaced and retained at 4 °C while particles were incubated in 150 µL PBS at 50 °C for 2 further weeks to fully degrade all NVP and CEA particles. Primary Human Skeletal Muscle Cells (SkMC; PromoCell) were cultured in SkMC media (PromoCell) and seeded at 16k cells/well in a 96-well plate overnight at 37 °C, 5% CO_2_. Media was exchanged to 90 µL/well, particle degradation buffer or fresh PBS was dosed at 10 µL/well in three technical replicates per batch, and incubated for 24 hours at 37 °C, 5% CO_2_. Treatment media was removed, analysed for acute cytotoxicity via the LDH-Glo assay (ProMega) according to the manufacturer’s instructions, and replaced with 1X alamarBlue reagent (Invitrogen) in SkMC media. After 24 hours further incubation, cell viability was determined by reading fluorescence at 570 nm excitation/585 nm emission. Relative cytotoxicity and viability were calculated relative to buffer-only controls.

### Hollow microparticle fabrication

Particles were loaded by applying vacuum after submersion in aqueous cargo solution as in previous work^4,26–28^ using 75 µL of 20% w/v CF647-OVA dissolved in 50% glycerol water. After air bubbles were visibly removed from the particle core, vacuum was released, and excess cargo was removed by pipette and Kim Wipe. Particles were then washed with ultrapure water by gently pipetting to remove surface-adsorbed protein and air dried.

Particles were sealed utilizing a modified dip-sealing technique,^27^ substituting a UV-curable PLGA-TA based resin for thermosensitive polycaprolactone used in prior work. In brief, particle arrays were fixed onto the top plate of a rheometer (Anton Paar), and slowly lowered into transient contact with a thin layer of PLGA-TA sealant, which was subsequently bonded to the particle lid features under UV light for 2 hours. The sealing layer substrate was prepared by spreading 75 µL of sealing resin in DCM (see above) on a glass slide to match the area of the particle array, and spin-coated at 2000 rpm for 15 s before being secured onto the bottom plate of the rheometer for dip-sealing. A single sealing layer substrate was suitable for sealing up > 10 particle arrays in series and could be monitored by observing particle indentations in sealant by eye as well as the deposition of sealant on particle lids by brightfield microscopy.

### Protein loading quantification

Particles of each chemistry (usually 15-30 particles) were counted and mechanically disrupted in a 1 mL Dounce homogenizer with 300-500 µL PBS. The debris was sedimented *via* centrifugation and 150 µL supernatant was analysed *via* microBCA (Thermo Fisher) according to the manufacturer’s instructions to determine protein concentration and thus loading/particle.

### *In vitro* protein release and particle degradation

Loaded and sealed particle batches (N = 4-5; 49 particles/batch) of each resin chemistry were immersed in PBS and gently detached from the substrate with a glass Pasteur pipette. Particles were incubated at 37 °C overnight and then washed with PBS three times to ensure full removal of surface absorbed protein and from any imperfectly sealed particles. Particles were then incubated in 300 µL PBS/batch at 37 °C, with 150 µL PBS replaced at indicated timepoints and stored at 4 °C for subsequent protein quantification by microBCA. Percent protein release was normalized to the full release per batch for CEA and NVP samples (which were macroscopically degraded) or to the average loading of separately measured DEGDA batches (which were not macroscopically degraded at study end).

In parallel with release studies, particles were imaged with a brightfield microscope (EVOS XL Core; 20X objective) at indicated timepoints until experiment conclusion at week 10. For each timepoint, 10 separate particles were measured in ImageJ. Particle swelling was defined as the distance from the two most distant points along the major axis relative to the starting length.

Scanning electron microscopy was performed using the methods above on unloaded, unsealed particles which had been incubated in PBS at 37 °C for the timepoints specified.

### *In vitro* antigen stability

Three batches of 25 µm thick, 9% NVP (w/w) particles were loaded with OVA and sealed before incubation in 200 µL PBS at 37 °C for one month. OVA was extracted by Dounce homogenization, concentrated on a 3 kDa-cutoff Amicon spin filter, and compared to fresh OVA using the Agilent Protein 230 kit on a 2100 Bioanalyzer instrument according to the manufacturer’s instructions.

### *In vivo* release and immunogenicity

All animals were handled in accordance with the UK Home Office Animals Scientific Procedures Act 1986 and with an internal ethics board and UK government approved project (PP5168779; Tregoning) and personal license (I87985646; Peeler). Food and water were supplied ad libitum. Female BALB/c mice (Charles River, UK) 6−8 weeks of age were placed into groups (N = 5) and housed in an acclimatized room.

To determine antigen dosing/particle, particle batches were pooled (usually 5-6 batches) for each chemistry and loading was quantified using a subset of particles as described for *in vitro* studies. Prior to injection, loaded and sealed particles were sterilized by immersion in ethanol before suspension in sterile PBS + 3% carboxymethylcellulose (w/v; viscosity enhancer) as in previous work.^3,4^ Soluble CF647-OVA and unlabelled OVA were likewise dissolved in sterile PBS + 3% methylcellulose (w/v) to achieve the desired dose. Mice were maintained under anaesthesia in a nose cone apparatus using isoflurane to ensure injection consistency. The neck scruff area was shaved to reduce fluorescent background and sterilized with ethanol before injection. Particle/antigen injection in the subcutaneous neck scruff was performed using 21-gauge needles to deliver 200 µL per mouse. After withdrawal, needles were flushed with PBS to confirm that all particles were injected.

At each imaging timepoint (twice weekly), mice were re-shaved as needed, anesthetized, and imaged with a Spectral Instruments Ami imaging system (N = 3-5 per group). Fluorescent imaging was performed with an exposure time of 10 seconds with the following parameters: 75% LED power, excitation 640 nm, emission 690 nm, binning 8, F Stop 2, object height 1.5 cm. Two images were taken per group to serve as technical replicates, with mice re-arranged in the chamber to randomize sensor positioning per mouse. In some cases, injections were inadvertently too deep for light penetration; these mice were excluded from imaging but not immunogenicity analysis.

Imaging data were analysed in Spectral Instruments Aura software (version 4.0.7) to report the fluorescent efficiency in photons/sec (angle-normalized radiance) from a uniform region of interest (ROI) at the injection site. The same size/shape ROI was used for each mouse and background was subtracted using an off-target ROI in each image. Background-subtracted fluorescent efficiencies were normalized to the maximum signal observed for that mouse over the course of the study to account for differences in injection depth and plotted as % maximum efficiency. The rate of cargo release was taken as the rate of fluorescent efficiency decay at the injection site and fitted with the non-linear least squares model “plateau followed by one phase decay” (Prism GraphPad) to account for particle hydration prior to release. Three mice were analysed per particle treatment and five mice were analysed per soluble protein treatment.

### Serum antibody concentration quantification

Mouse blood was collected via tail vein puncture at weeks 2, 4, 6, and 8 after injection, allowed to clot overnight in a microcentrifuge tube, and centrifuged at 5000 g for 20 minutes at 4 °C to yield serum in the supernatant. Anti-OVA IgG antibody titers were measure using an end point ELISA. High-binding MaxiSorp plates (Thermo Fisher) were coated with OVA at 10 μg/mL (for samples) or a 1:1 mixture of anti-mouse kappa and anti-mouse lambda capture antibodies at 5 µg/mL (for standards; Southern Biotech) in PBS at 4 °C overnight. Highly purified polyclonal mouse IgG, IgG1, IgG2a, or IgG2b (Southern Biotech) were used to generate a standard curve in a 1:5 dilution down the plate, starting at 200 ng/ml. Plates were washed three times and then blocked with assay buffer (PBS + 1% BSA [w/v] + 0.05% Tween-20 [v/v]) for 1 h at 37 °C. Serum samples were serially diluted in assay buffer and incubated on the coated plates for 2 h at 37 °C; 10000-fold diluted serum was used for most assays, with the exception of 1000-fold diluted serum used to ensure sensitive detection of IgG subtype titers. Plates were washed three times with assay buffer, and HRP-conjugated donkey-anti-mouse IgG, IgG1, IgG2a, or IgG2b (1:10000, Southern Biotech) was added for 1 h at 37 °C. Plates were washed three times with assay buffer, developed with SureBlue TMB Microwell Peroxidase Substrate and stopped with 2 N H_2_SO_4_ (Insight Biotechnology Ltd). After development, plates were read at 450 nm using a VersaMax plate reader. Generation of the standard curve and anti-OVA IgG quantification in samples was performed following subtraction of background optical density (OD) values, followed by interpolation of serum concentrations using a sigmoidal, four point least squares fit of the standard curve (GraphPad Prism).

### *In vivo* biocompatibility

The skin surrounding the neck scruff injection site was dissected after sacrifice and pinned to a cork dissection board for examination of the subcutaneous facia for inflammation. Tissue was fixed with PBS + 4% paraformaldehyde (v/v) for 1 h, photographed, and stored in OCT compound at -80 °C. Samples were later washed with PBS to remove OCT and imaged with a MRCL700 3D Imager Pro (Microqubic AG) and associated viewing software.

## Statistical Analysis

All results are expressed as a mean ± standard deviation (SD) as indicated in the Fig. captions. One-way ANOVA test with a Tukey’s multiple-comparisons test was used for comparison across multiple groups. Statistical analysis and curve fitting were performed in Prism 10.3.1 (GraphPad Software). Statistical significance was considered as p < 0.05.

## Acknowledgements

This work was supported in part by the Bill & Melinda Gates Foundation [INV-003863]. The conclusions and opinions expressed in this work are those of the author(s) alone and shall not be attributed to the foundation. Under the grant conditions of the foundation, a Creative Commons Attribution 4.0 License has already been assigned to the Author Accepted Manuscript version that might arise from this submission. Please note works submitted as a preprint have not undergone a peer review process.

D.J.P., J.Y., O.R.-G., and V.L. were supported by the European Union’s Horizon 2020 research and innovation programme under Marie Skłodowska-Curie grant agreements 101027174, 839137, 893158, and 890854, respectively. M.M.S. and J.P.W. acknowledge funding from the grant from the UK Regenerative Medicine Platform “Acellular / Smart Materials – 3D Architecture” (MR/R015651/1). M.M.S. acknowledges support from the Royal Academy of Engineering Chair in Emerging Technologies award (CiET2021\94) and the Rosetrees Trust.

The authors thank Paul McKay, Karnyart Samnuan, Luke Granger, Ziyin Wang, Hadijatou Sallah, Chubicka Thomas, and the rest of the Group of Mucosal Immunology for invaluable help with reagent sourcing and immunological techniques. We are further grateful to Imperial College London Central Biomedical Services for assistance with animal work, especially Kasia Smigielska, Mark Harrington, Amy Wathen, and Justyna Glegola. Lastly, we are grateful to Dennis Lee and Jeremy Blum for their guidance throughout this project.

## Data availability

All the data supporting the results in this study are available within the paper and its Supplementary Information. Source data are provided with this paper. Additional data may be requested from the authors.

## Contributions

The grant funding this work was conceived by J.Y., O.R.-G., J.P.W., R.J.S. and M.M.S. M.M.S. contributed to project design and supervised the project. Macromonomer synthesis and purification methods were developed by J.Y., O.R.-G., J.P.W., D.J.P. and C.K. Resin formulation and printing was developed by D.J.P., K.Z. and R.S. In vitro experiments were prepared and carried out by D.J.P., R.S., C.K., P.P., K.Z., G.B., J.P.W., T.F.D., V.L., X.S., and K.P. In vivo experiments were performed by D.J.P. and J.S.T. with support from R.S. and P.P. Data analysis was performed by D.J.P., R.S., C.K., J.P.W. and V.L. The paper was written by D.J.P. with input from all co-authors.

## Corresponding authors

Correspondence to Molly M Stevens.

