## Supplemental Figures for "“Nanoscale biodegradable printing for designed tuneability of vaccine delivery kinetics”"

1. PLGA-TA characterization by $^{1}$H NMR
2. PLGA-TA characterization by $^{13}$C NMR
3. PLGA-TA characterization by LC-MS
4. PCL-TA characterization by $^{1}$H NMR
5. PCL-TA vs PLGA-TA print characterization by SEM
6. Print resolution vs laser power SEM for various co-monomers
7. Print resolution vs laser power SEM for down-selected co-monomer blends
8. AT-FTIR analysis of vinyl conversion
9. Particle design evolution
10. Quantification of loading for each resin chemistry
11. Co-monomer dependent particle degradation at 50 °C
12. Encapsulated antigen stability by bioanalyzer
13. Release quantification for prime-boost study
14. Antibody subtyping of week 8 serum from prime-boost immunization
15. Subcutaneous neck scruff biocompatibility
16. Subcutaneous flank biocompatibility

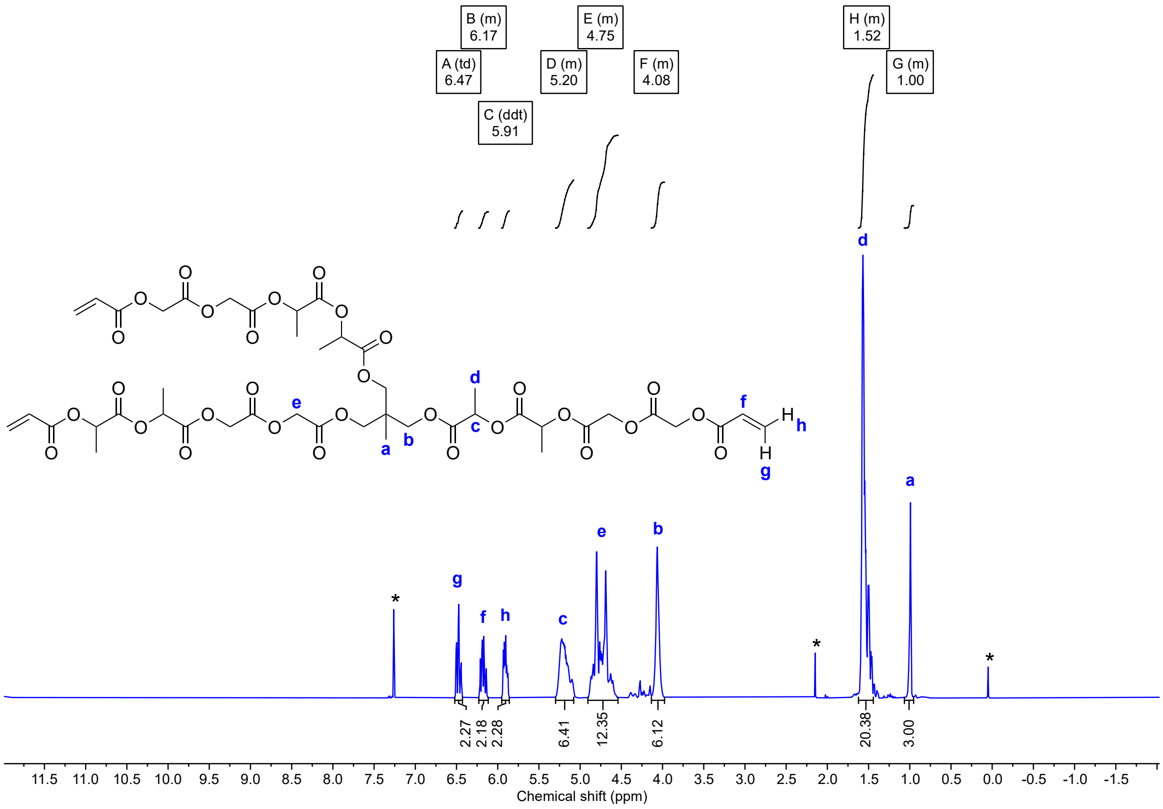

**Figure S1**. $^{1}H$ NMR spectra of PLGA-TA in $CDCl_3$ at 500 MHz. The chemical structure shown represents the average chemical structure of one possible isomer (poly($L_6GA_6$)-triacrylate). Comparison of the integrations of the vinyl protons ($H_f$, $H_g$, $H_h$) (δ = 6.47 – 5.91 ppm) referenced to the initiator $CH_3$ protons ($H_a$) indicates a degree of functionalisation of 75%. * indicates residual solvent (δ = 7.26 ppm), acetone (δ = 2.17 ppm) and tetramethylsilane (δ = 0 ppm).

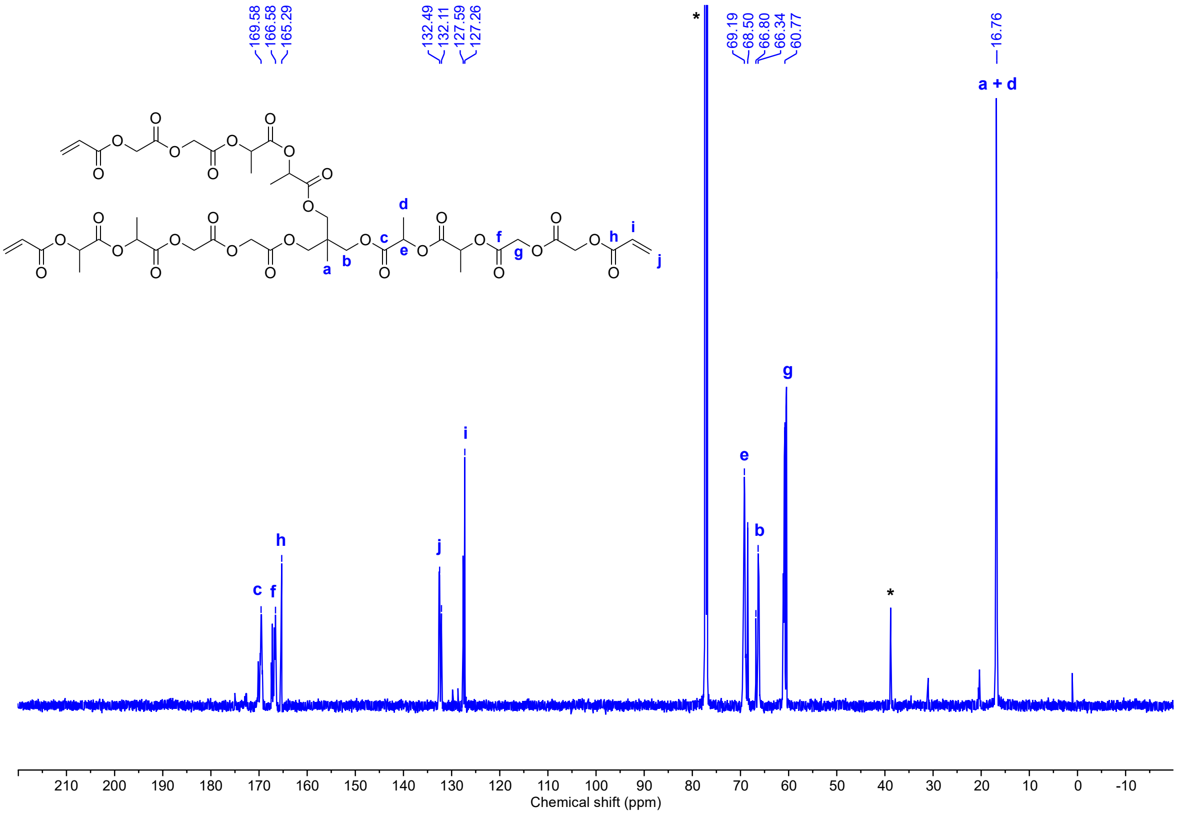

**Figure S2**. $^{13}C$ NMR spectra of PLGA-TA in $CDCl_3$ at 125 MHz. The chemical structure shown represents the average chemical structure of one possible isomer (poly($L_6GA_6$)-triacrylate). * indicates residual solvent (δ = 77.16 ppm), acetone (δ = 30.92 ppm) and tetramethylsilane (δ = 0 ppm).

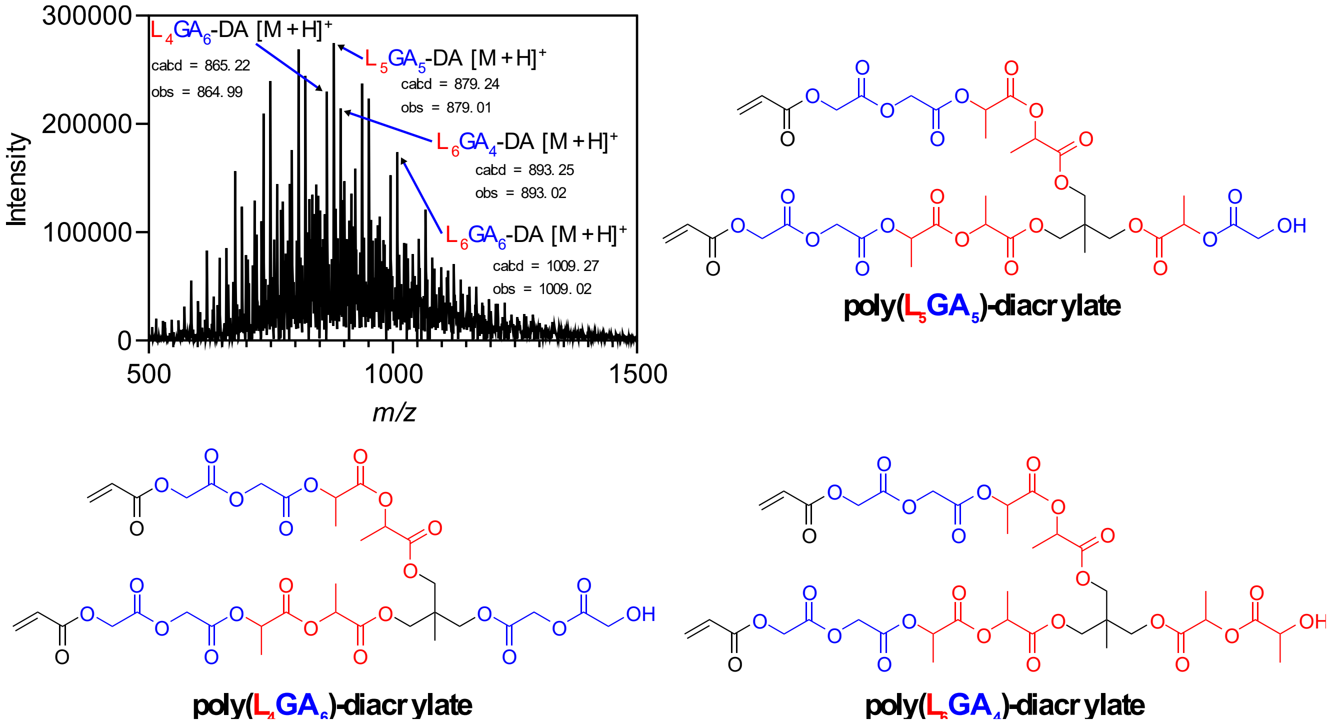


**Figure S3**. Positive mode ESI/APCI mass spectrum of PLGA-TA extracted from the centre of the polymer envelope (4.502 - 5.001 min). The dominant structure identified in the ESI/APCI mass spectra corresponds to a diacrylate isomer with a degree of polymerisation corresponding to five lactic acid and five glycolic acid monomers (poly($L_5GA_5$)-diacrylate). Whilst the mass spectrum shows the poly($L_5GA_5$)-diacrylate as the dominant polymer, based on NMR and GPC analysis, the main component in the mixture is the poly($L_6GA_6$)-triacrylate polymer. The dominance of the diacrylate derivative in the mass spectrum is most likely to be due to the higher probability of observing this mass adduct due to protonation of the terminal hydroxyl group. This result agrees with the NMR characterisation which demonstrates an average degree of functionalisation of 75% of the hydroxyl groups on the three-arm PLGA polymer. It is speculated the odd numbered incorporation of lactic acid and glycolic acid units (i.e. Dp = 5 each) is a result of the synthesis conditions, which utilises high temperatures (130 °C) in the presence of a tin(II) ethylhexanoate catalyst which has been reported in related derivatives to give 3-methylglycolides.

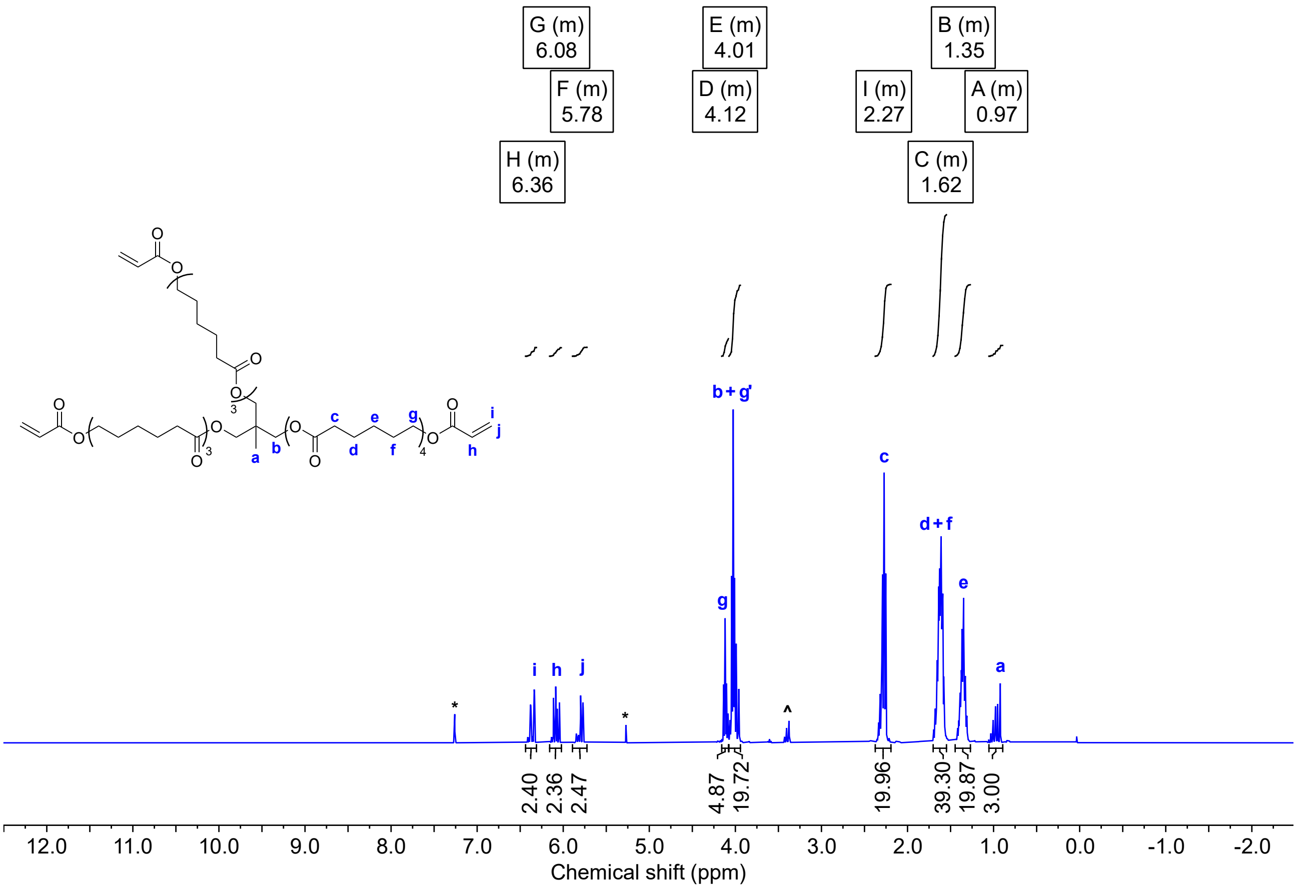


**Figure S4**. ¹H NMR spectra of PCL-TA in $CDCl_3$ at 400 MHz. The chemical structure shown represents the average chemical structure of one possible isomer (poly($CL_{10}$)-triacrylate). Comparison of the integrations of the vinyl protons ($H_h$, $H_i$, $H_j$) ($\delta$ = 6.36 – 5.78 ppm) referenced to the initiator $CH_3$ protons ($H_a$) indicates a degree of functionalisation of 80%. Protons $H_{g'}$ (d = 4.01 ppm) correspond to the α-$CH_2$ protons. ^ Indicates suspected $H_g$ protons corresponding to unfunctionalised alcohol end groups. * Indicates residual solvent ($\delta$ = 7.26 ppm), dichloromethane ($\delta$ = 5.30 ppm).

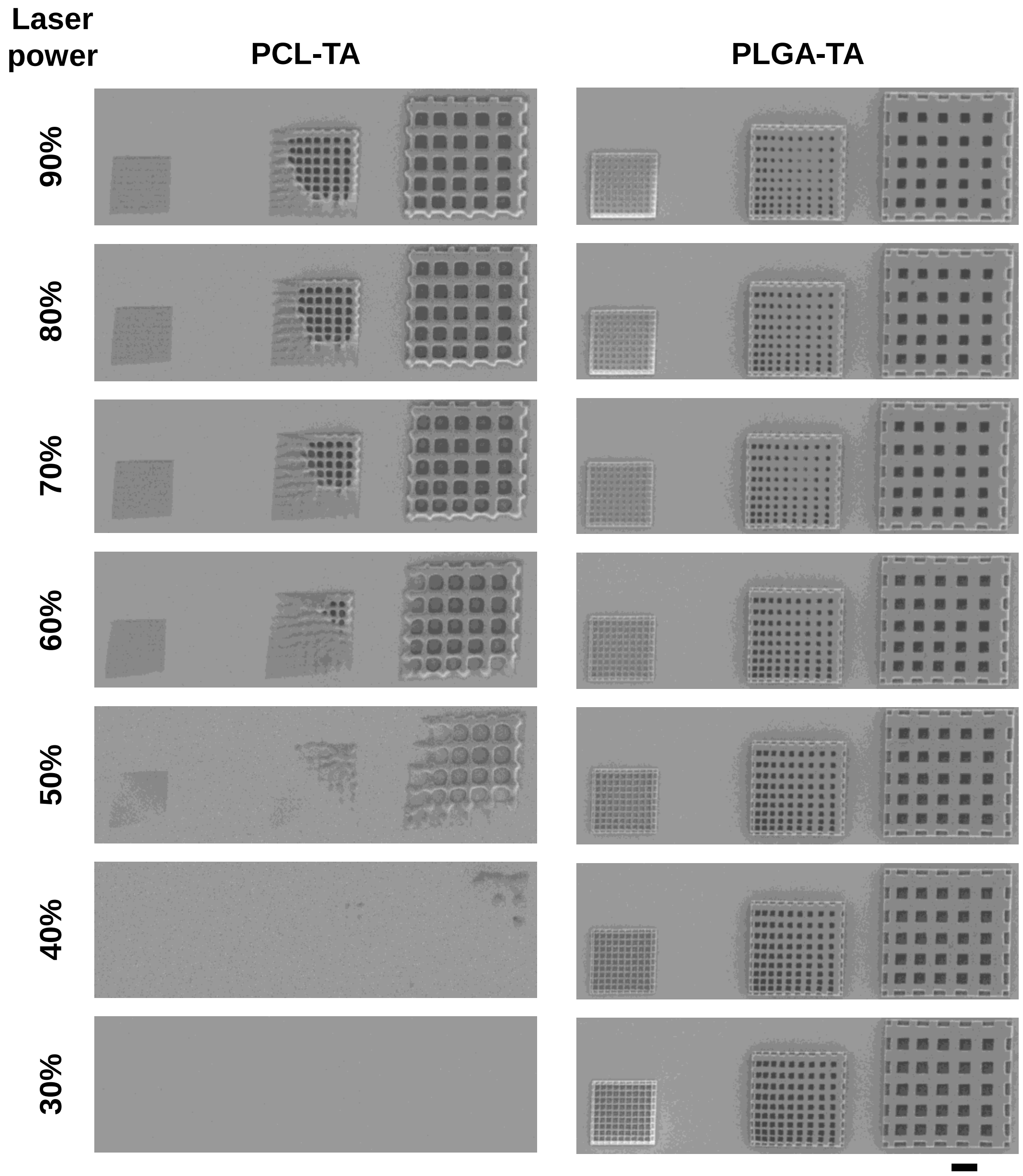


**Figure S5**. Poly(lactic-co-glycolic acid) triacrylate has higher polymerization reactivity than poly(caprolactone) triacrylate. Grids were printed with triacrylate resin composed of 19% NVP + 1% w/w EMK at the laser powers shown and imaged with scanning electron microscopy at 600X magnification. Scale bar = 20 µm.

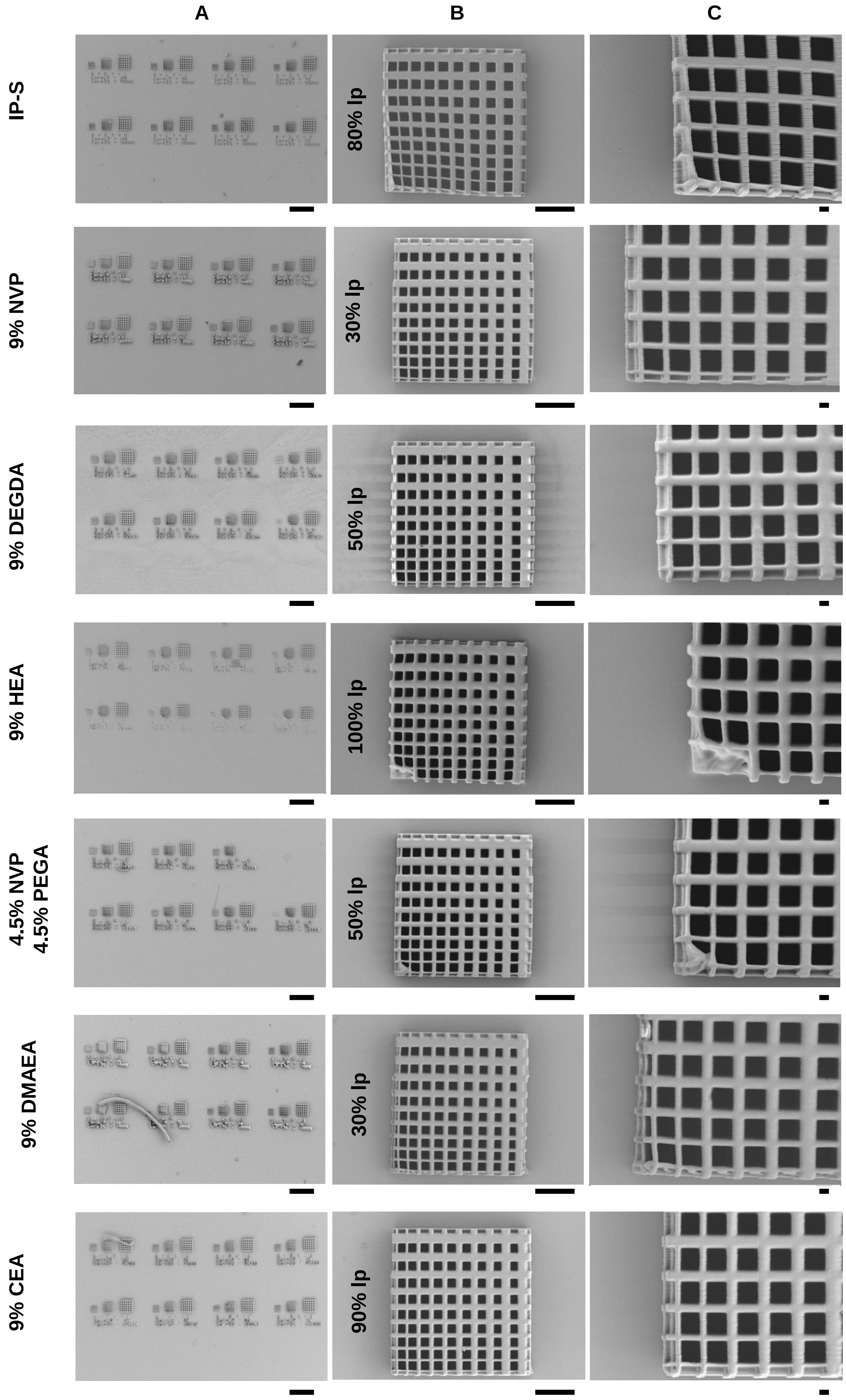


**Figure S6**. Laser power and scan speed optimization towards maximum resolution and efficiency. Grids were printed with varying laser power (ranging from 20-100% in 10% increments) and scan speed (ranging from $10^4$ to $10^5$ mm/s in $10^4$ increments) using either commercial IP-S resin or PLGA-TA + 1% w/w EMK + the indicated weight percent of various comonomers and imaged with scanning electron microscopy. (A) Subset of the laser power vs. scan speed optimization; scale bar = 200 µm. (B) Optimal laser power $10^5$ mm/s scan speed; scale bar = 20 µm, and (C) Closer view of optimal laser power $10^5$ mm/s scan speed; scale bar = 2 µm.

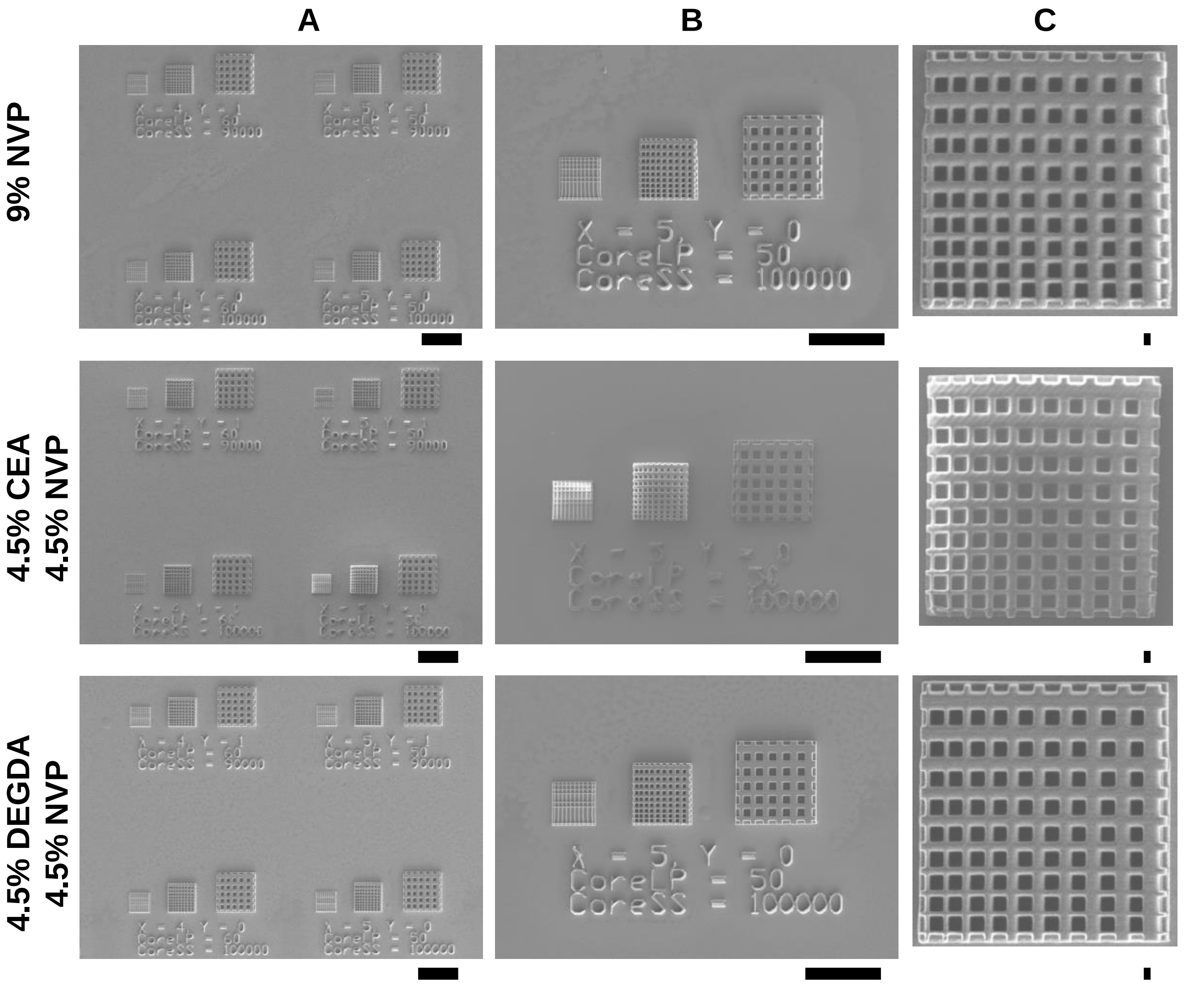


**Figure S7**. Laser power and scan speed optimization for NVP/comonomer blend resins. Grids were printed with varying laser power (ranging from 20-100% in 10% increments) and scan speed (ranging from $10^4$ to $10^5$ mm/s in $10^4$ increments) using PLGA-TA + 1% w/w EMK + the indicated weight percent of various comonomers and imaged with scanning electron microscopy. (A) Subset of the laser power vs. scan speed optimization array; scale bar = 100 µm. (B) 50% laser power grids at $10^5$ mm/s scan speed; scale bar = 100 µm. (C) Closer view of the 50% laser power at $10^5$ mm/s scan speed grid; scale bar = 1 µm.

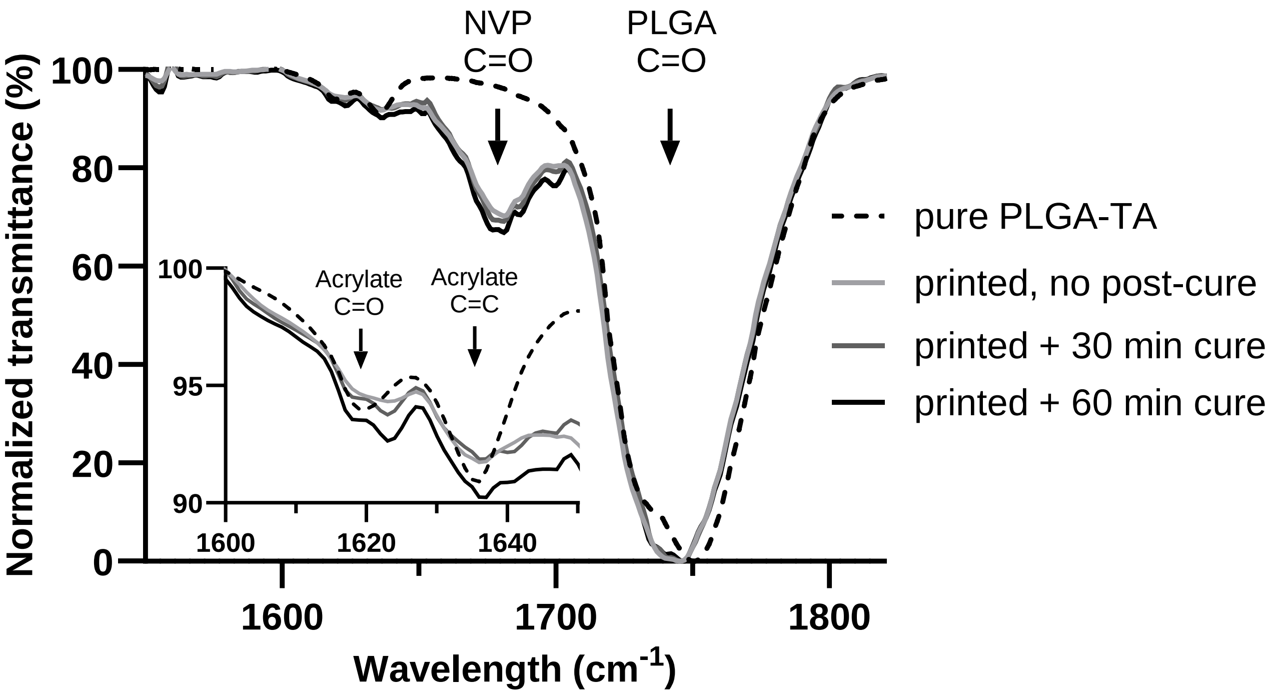


**Figure S8**. AT-FTIR spectra of pure PLGA-TA and NVP SCRIBE resin printed at 80% laser power and post-cured with UV light for varying times. Data are plotted as the mean of 52 scans.

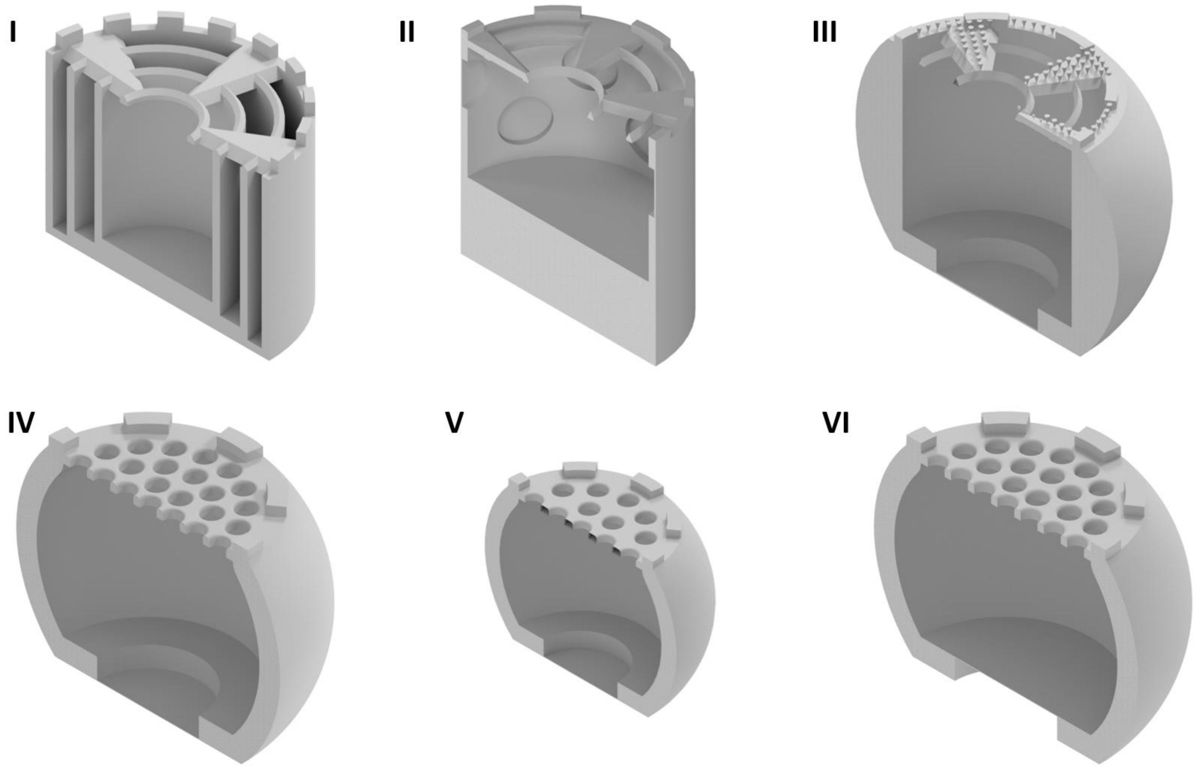


**Figure S9**. Design evolution towards CAD-controlled release. I) Multiple chambers within one microparticle; II) Side portholes with tunable side thicknesses; III) Bottom porthole with bumpy lid to enhance sealing adhesion; IV) Bottom porthole with refined lid for more efficient loading, cleaning and sealing; V) Bottom porthole with reduced volume for easier injection; VI) Raised bottom porthole for more reliable thickness-controlled drug release (final design).

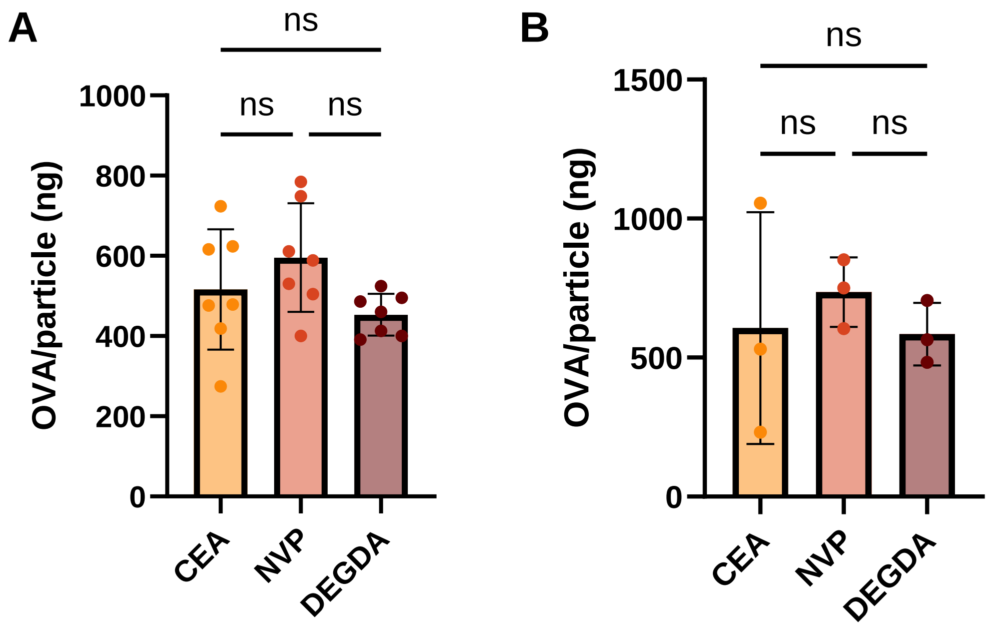


**Figure S10**. Particle loading is not dependent on porthole membrane thickness or resin chemistry. A glass homogenizer was used to mechanically disrupt OVA-loaded particles with porthole thickness of either (A) 1 µm or (B) 25 µm. After disruption, protein content in the supernatant was analyzed by microBCA and normalized to particle number. The data are presented as the mean ± SD (N = 3-7 batches with 49 particles each). As detailed in the methods section, particles batches were pooled and redistributed before injection *in vivo*.

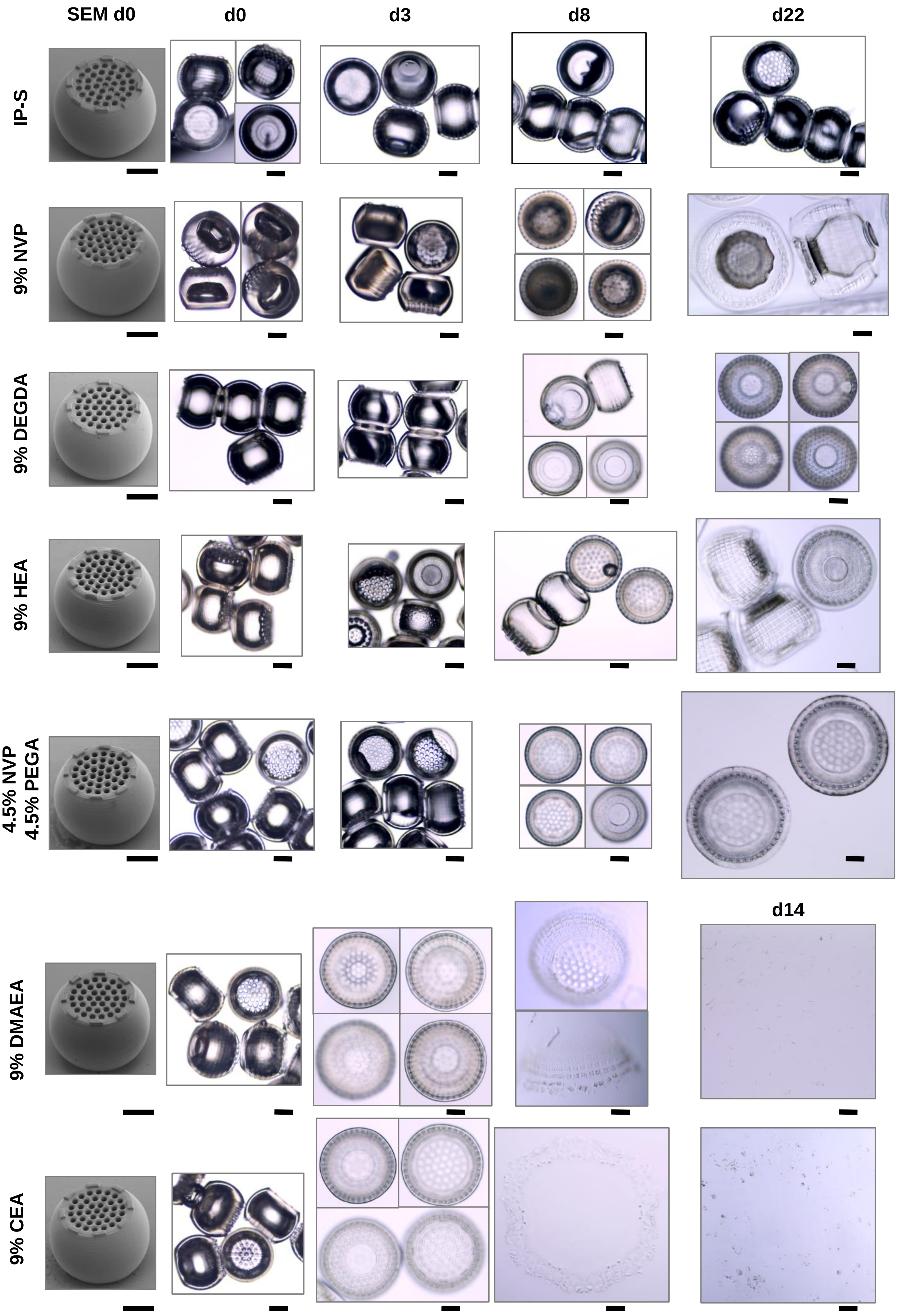


**Figure S11**. Comonomer chemistry dictates particle swelling and degradation rate. Particles were printed with either commercial IP-S resin or PLGA-TA + 1% w/w EMK + the indicated weight percent of various comonomers at the optimal laser power determined from the resolution grid. Particles were either imaged with scanning electron microscopy or incubated at 50 °C in PBS and imaged with 20X light microscopy images at the times indicated. Scale bar = 100 µm. Individual cropped images have been outlined to enable juxtaposition of particles from the same sample and time point.

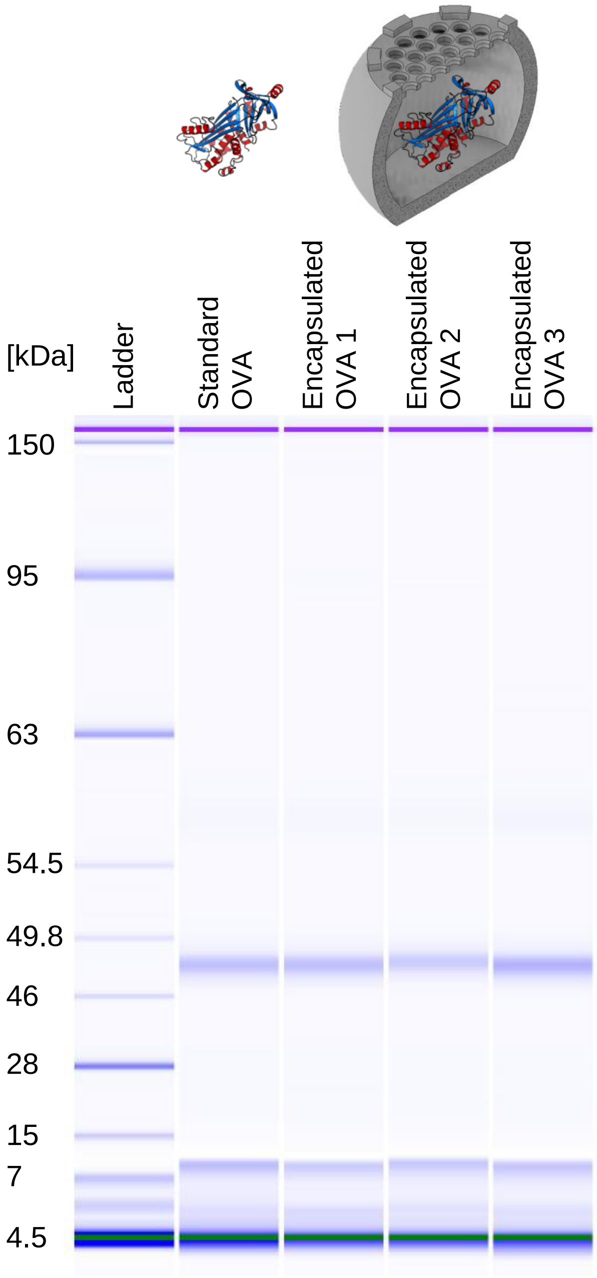


**Figure S12**. Characterizing particle-loaded OVA stability with gel electrophoresis. OVA was loaded in 50% glycerol into 25 µm-thick 9% NVP particles, incubated for 1 month at 37 °C in PBS, and then retrieved by mechanical disruption for molecular weight analysis using the Bioanalyzer Protein 230 kit (N=3 batches of 49 particles). The molecular weight of OVA extracted from encapsulated particles matches the single 48 kDa band observed for the control, non-incubated protein.

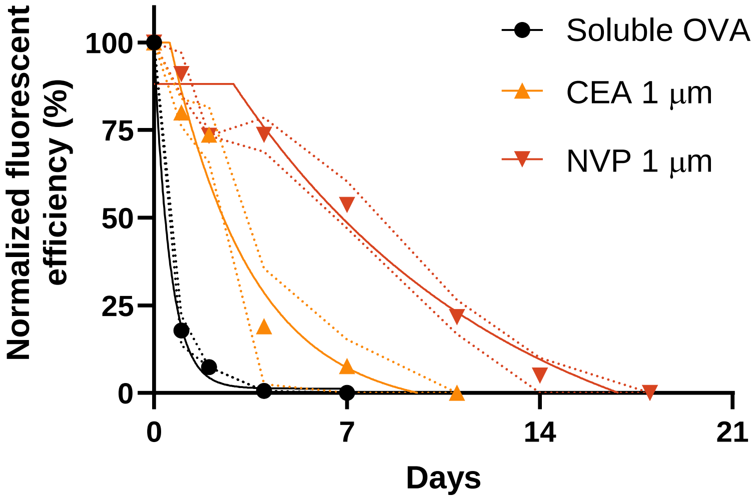


**Figure S13**. Longitudinal *in vivo* fluorescent imaging of BALB/c mice injected subcutaneously with either 5 µg soluble CF647-OVA or 5 µg soluble OVA + 20 µg CF647-OVA encapsulated in CEA or NVP particles with 1 µm porthole thickness. Background-subtracted injection site ROI fluorescent efficiency ($\lambda_{ex}/\lambda_{em}$ = 640/690 nm) was normalized to the maximum observed per mouse (Data shown as mean [circles] ± SD [dashed lines] with the plateau followed by one phase decay fit [solid line; N=3-4]). Note that a different batch of CF647-OVA was used for this study (1:10 CF647:OVA labelling) compared to the prime-only study (1:1 CF647:OVA labelling), leading to a faster apparent decay of signal due to the limit of detection.

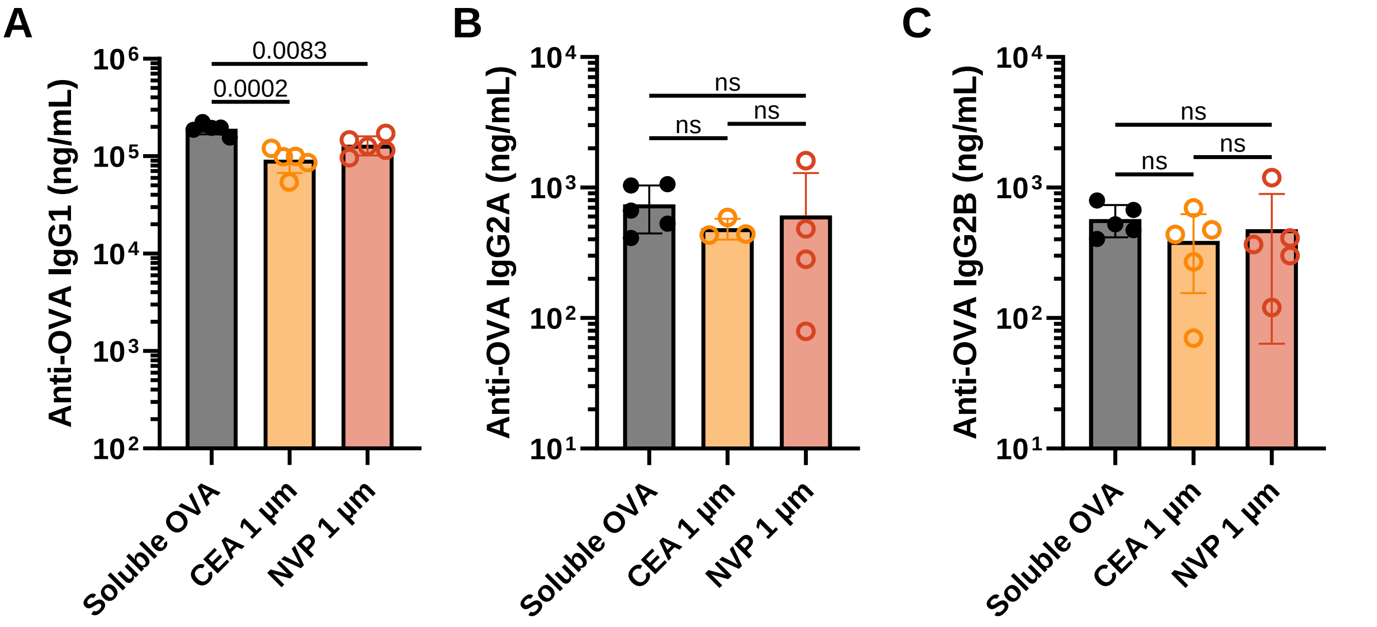


**Figure S14**. Antibody subtyping of week 8 serum from prime-boost immunization. Anti-OVA (A) IgG1 titers, (B) IgG2A titers, (C) IgG2B titers plotted as mean ± SD (N=3-5; p-values via one-way ANOVA with Tukey's multiple comparisons test).

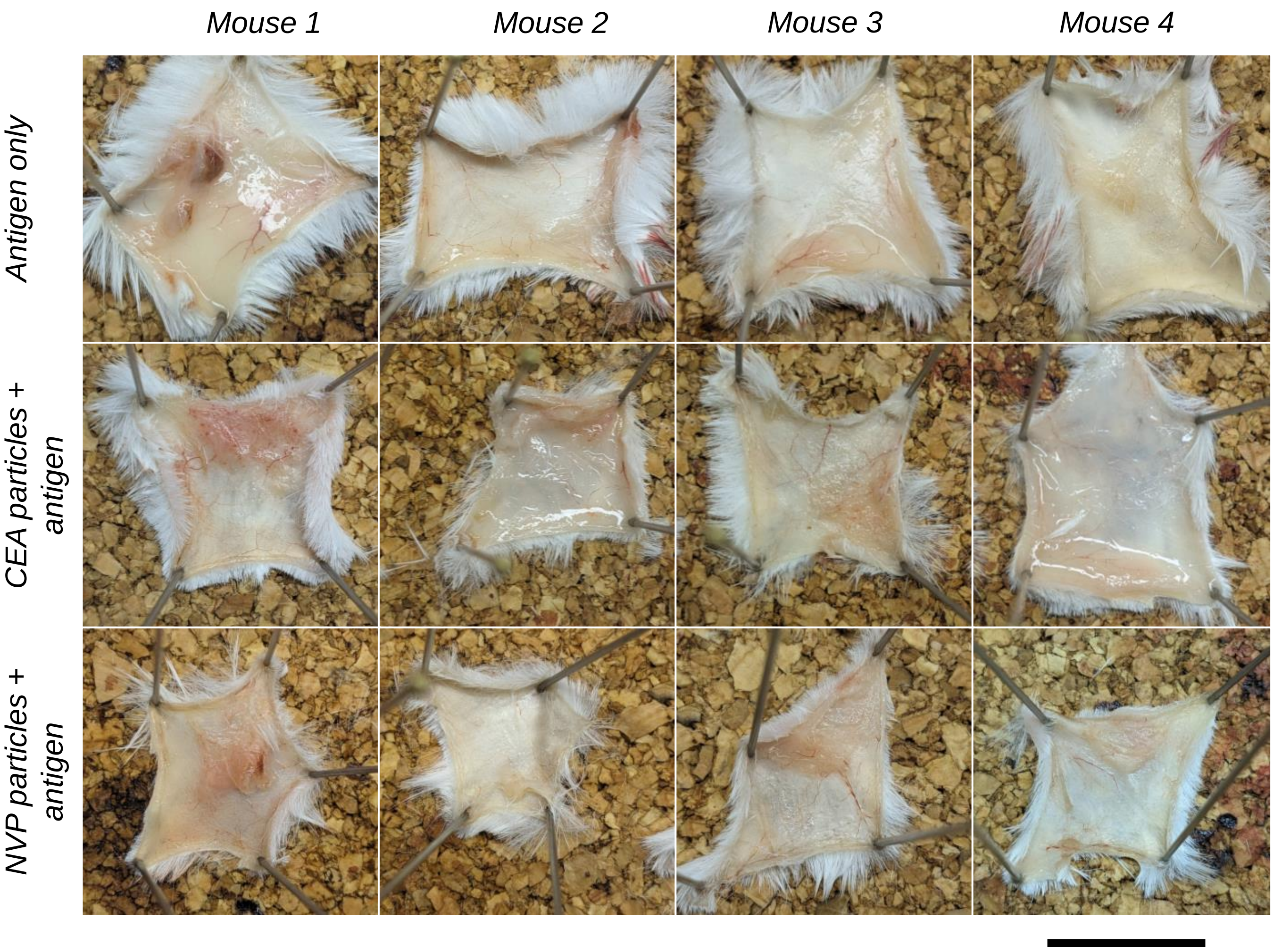


**Figure S15**. Dissected subcutaneous neck scruffs at day 61 demonstrate particle biocompatibility. Mice were sacrificed 61 days after subcutaneous injection of soluble or particle-encapsulated antigen, and the injection site was dissected, pinned, and fixed with PBS + 4% paraformaldehyde before imaging. No difference in redness or swelling indicative of unresolved inflammatory processes were discerned between groups. Scale bar = 2 cm. Individual cropped images have been outlined in white.

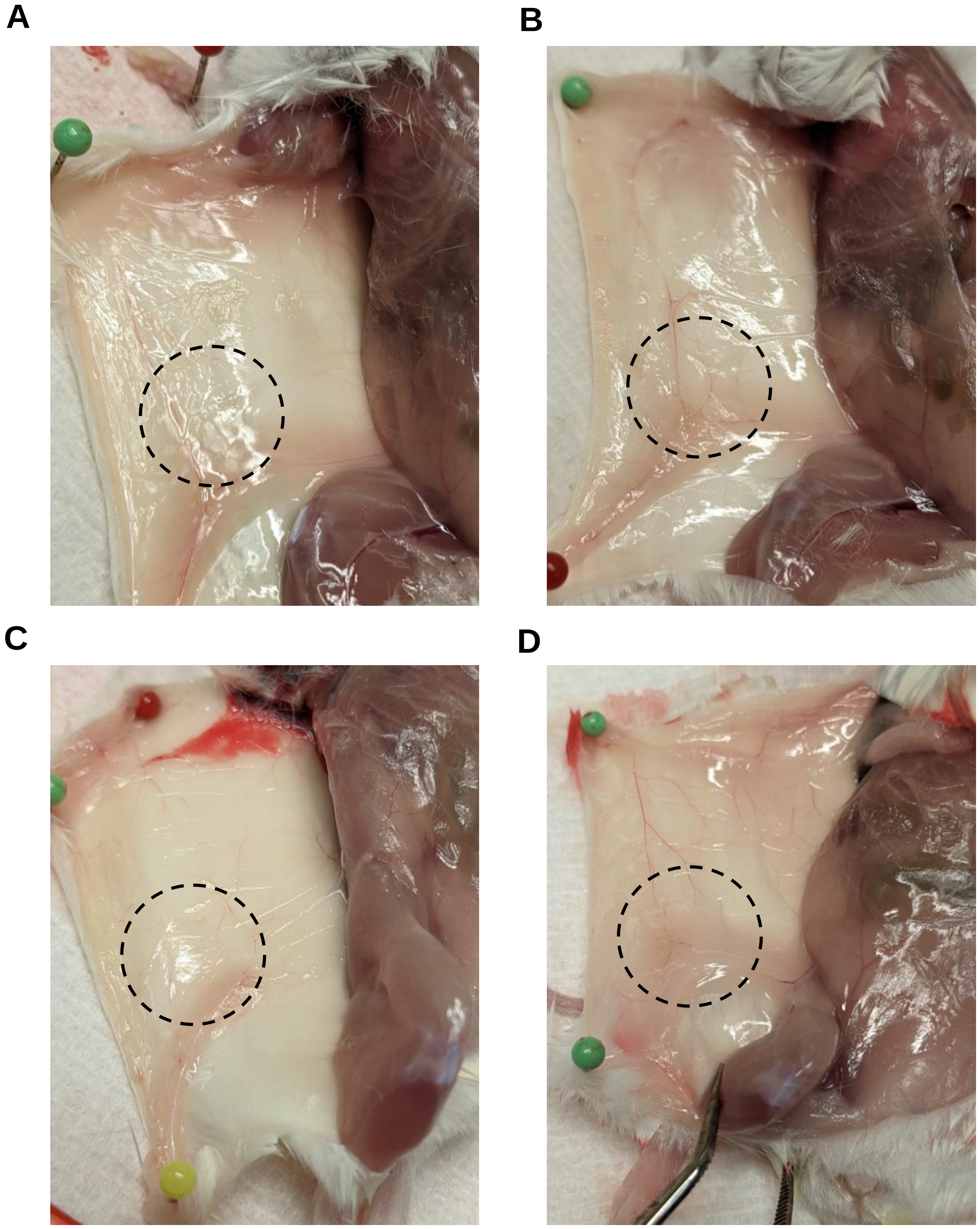


**Figure S16**. Dissected subcutaneous flanks at day 49 demonstrate particle biocompatibility. Mice were sacrificed 49 days after subcutaneous injection of soluble (A) or particle-encapsulated (B-D) dextran, and the injection site (dashed circle) was exposed before imaging. No difference in redness or swelling indicative of unresolved inflammatory processes were discerned between groups.
